# Pln1 Mediates Lipid Droplet-Vacuole Tethering During Microlipophagy in *Saccharomyces cerevisiae*

**DOI:** 10.1101/2025.05.07.652694

**Authors:** Monala Jayaprakash Rao, Brayden Folger, Joshua Reus, Alexandre Toulmay, Jin Li, Rudian Zhang, William Prinz, Joel Goodman, Fei Wang

**Affiliations:** Department of Cell Biology, UTSW; Department of Pharmacology, UTSW

## Abstract

Lipid droplets (LDs) are dynamic organelles that undergo growth or degradation depending on the metabolic state of the cell. One form of LD degradation is autophagy-mediated, referred to as lipophagy. Here, we demonstrate that Pln1, a perilipin located on the surface of LDs in *Saccharomyces cerevisiae*, previously known for its role in LD biogenesis, is essential for lipophagy. Pln1 facilitates the docking of cytosolic LDs to vacuoles, the lysosome-like organelles responsible for LD degradation, under various nutrient conditions. Molecular dissection of Pln1 revealed that the N-terminal PAT (Perilipin (PLN1), Adipophilin (PLN2), and TIP47 (PLN3)) domain and a hydrophobic region are critical for the localization and binding of LDs to vacuoles. Site-specific mutagenesis within the PAT domain identified a semi-hydrophobic LD Interacting Motif (LIM), which is vital for this interaction. Furthermore, an intrinsically disordered region (IDR) near the center of Pln1 is required for efficient LD-vacuole tethering. These findings support a model in which Pln1 bridges LDs and vacuoles by simultaneously interacting with both organelles. Notably, deleting PLN1 did not impair survival during prolonged nitrogen starvation and enhanced viability in autophagy-defective (*atg8*Δ) cells, suggesting that balancing Pln1-mediated LD biogenesis and lipophagy is crucial for yeast survival under starvation conditions.

## Introduction

Lipid droplets (LDs) are specialized organelles found in all eukaryotic cells, primarily serving as storage depots for neutral lipids, such as triglycerides (TAG) and steryl esters (SE). These organelles originate as extensions from the cytosolic leaflet of the endoplasmic reticulum (ER) and are characterized by a core enriched with TAG and/or SE, surrounded by a phospholipid monolayer embedded with proteins that are critical for LD metabolism and stability (Athenstaedt & Daum, 2006). In *Saccharomyces cerevisiae*, the enzymes Dga1 and Lro1 contribute to TAG synthesis from diacylglycerol (DAG), while SE synthesis occurs through the acylation of sterols by Are1 and Are2 (Jensen-Pergakes et al., 2001; Oelkers et al., 2002; Oelkers et al., 2000). Proteins such as seipin, Pln1, and Fit2 facilitate LD formation and organization, working in concert with other lipid metabolic enzymes, structural components, and regulatory proteins (Arlt et al., 2022; Cartwright et al., 2015; Gao et al., 2017; Rao & Goodman, 2021). Across species, from yeast to humans, the complex interplay between LD-associated proteins and lipids facilitates interactions between LDs and other intracellular structures, playing a role in various biological processes.

LDs undergo dynamic degradation in response to cellular metabolic demands (Diep et al., 2024; Olzmann & Carvalho, 2019; Paar et al., 2012). They act as crucial reservoirs for cellular energy storage, lipid buffers, and essential lipid components for signaling and membrane biogenesis (Farese & Walther, 2009; Moir et al., 2012; Rao et al., 2018). Dysregulation of LD degradation has been implicated in liver, neuronal, and metabolic diseases (Pressly et al., 2022).

LD lipids can be broken down through two primary pathways: cytosolic lipase-mediated lipolysis and autophagy-dependent LD degradation (lipophagy) (Schott et al., 2019; Zechner et al., 2017). Lipolysis involves well-characterized cytosolic lipases, including hormone-sensitive lipase (HSL) and adipose triglyceride lipase (ATGL) (in yeast, Tgl3, Tgl4, and Tgl5), which dock onto LDs to degrade lipids in the cytosol (Kurat et al., 2006). In contrast, lipophagy—similar to other forms of autophagy—can proceed via the formation of autophagosomes around LDs, which subsequently fuse with lysosomes (macrolipophagy), or through direct invagination of LDs by lysosomes (bypassing the autophagosome), followed by LD degradation by lysosomal lipases. In mammalian systems, macrolipophagy is supported by autophagic machinery, with lipophagy receptors such as spartin, Orp8, and ATG14 contributing to substrate selectivity (Chung et al., 2023; Pu et al., 2022; Yuan et al., 2024). These receptors recruit a subset of LDs to autophagosomes by interacting with ATG8/LC3 proteins via LC3-interacting region (LIR) motifs, although the mechanistic understanding of these receptors remains limited.

In contrast to macrolipophagy, the molecular machinery of microlipophagy is less understood, partially because macrolipophagy and other forms of bulk autophagy often overlap with microlipophagy. For example, general autophagy may contribute to the establishment or maintenance of membrane phase separation necessary for microlipophagy, possibly via sphingolipids (Yamagata et al., 2011). Additionally, some ATG proteins, such as Atg1 and Atg8 in yeast, are involved in all forms of autophagy, including microlipophagy (Nakatogawa et al., 2009). Despite these complexities, a key feature of macrolipophagy is the establishment of LD-lysosome (vacuole in yeast) contacts, which precedes lysosomal injection or engulfment of LDs. Recent findings related to LD-lysosome contacts include the identification of tether proteins such as Ldo16 and Ldo45 in yeast and ARL8B in macrophages (Álvarez-Guerra et al., 2024; Diep et al., 2024; Menon et al., 2023).

The machinery and regulation of lipophagy are context-dependent. In the liver, both macro- and microlipophagy are highly active (Schulze et al., 2020; Singh et al., 2009). In *Saccharomyces cerevisiae*, microlipophagy appears to be the primary pathway (Goodman, 2021; Schott et al., 2022; van Zutphen et al., 2014; Vevea et al., 2015), induced under various conditions. The factors responsible for triggering lipophagy do not entirely overlap between these conditions (Fairman & Ouimet, 2022). Recent reports show that during glucose starvation, the LD proteins Ldo16 and Ldo45 promote microlipophagy by tethering LDs to the vacuolar membrane through interactions with the vacuolar protein Vac8 (Álvarez-Guerra et al., 2024; Diep et al., 2024). However, it remains unclear whether Ldo16 and Ldo45 are involved in microlipophagy under other lipophagy-inducing conditions, such as nitrogen starvation. Furthermore, genetic deletion of both Ldo16 and Ldo45 only partially reduces LD-vacuole association and microlipophagy, indicating the presence of additional regulators involved in LD-vacuole contact formation during microlipophagy.

The perilipins (PLNs) are a family of proteins that bind, coat, and stabilize LDs in eukaryotic cells, regulating both lipolysis and lipophagy. For example, PLIN1 and PLIN5 suppress lipolysis by sequestering ABHD5, an ATGL co-activator, preventing its binding to ATGL in a phosphorylation-dependent manner. Conversely, degradation of PLIN2 and PLIN3 removes a barrier to both cytosolic lipolysis and autophagy. Thus, a common role of the PLNs is to regulate LD stability. Interestingly, yeast Pln1 supports TAG and LD biogenesis by stimulating Dga1 activity (Gao et al., 2017), a role not reported in mammalian cells. However, similar to mammalian PLNs, the yeast perilipin (Pln1) enhances LD integrity (Gao et al., 2017), although the underlying mechanism remains unclear.

In this study, we identified Pln1 as a positive regulator of lipophagy. Unlike the deletion of Atg1 (*atg1*Δ) or Atg8 (*atg8*Δ), deletion of Pln1 (*pln1*Δ) does not affect general autophagy but specifically inhibits lipophagy by reducing LD-vacuole association, suggesting that Pln1 is involved in microlipophagy. Remarkably, the lipophagy defect in *pln1*Δ cells was as severe as in *atg1*Δ or *atg8*Δ cells. Furthermore, Pln1 supports lipophagy during nitrogen starvation, glucose restriction, and the stationary phase, in contrast to Ldo16/Ldo45, which support microlipophagy during glucose starvation but not nitrogen starvation. Using truncation and site-specific mutagenesis, we identified regions and residues in Pln1 involved in its LD localization and in supporting LD-vacuole association. Finally, we demonstrated that genetic deletion of Pln1 (*pln1*Δ) did not impair cell survival during prolonged nitrogen starvation, suggesting that the dual function of Pln1 in LD biogenesis and turnover may partially offset the effects on cell survival. Consistent with this, *pln1*Δ improved cell survival in autophagy-defective cells (*atg8*Δ).

## Results

### Pln1 is Essential for Lipophagy but Not General Autophagy

Previous studies have established that Pln1 supports TAG synthesis (Gao et al., 2017). Although it has a role in LD biosynthesis (Gao et al., 2017), a possible role in LD degradation has not been investigated. In wild-type cells, but not in *pln1*Δ cells, Erg6-GFP, an LD-resident protein, was efficiently degraded in the vacuole under nitrogen starvation (SD-N) and glucose restriction (SD with 0.4% glucose), producing free vacuolar proteolysis-resistant GFP (Fig. 1A-B). The vacuolar degradation of GFP-tagged Erg6 (Erg6-GFP processing) is a common indicator of lipophagy (van Zutphen et al., 2014; Zhang et al., 2020). Remarkably, *pln1*Δ significantly reduced Erg6-GFP processing to levels similar to the deletion of Atg1, Atg8, Ypt7, and Pep4 (Fig. 1C, Fig. S1A-C), which are essential components for lipophagy. This suggests that Pln1 is a vital regulator of lipophagy.

**Figure 1.**
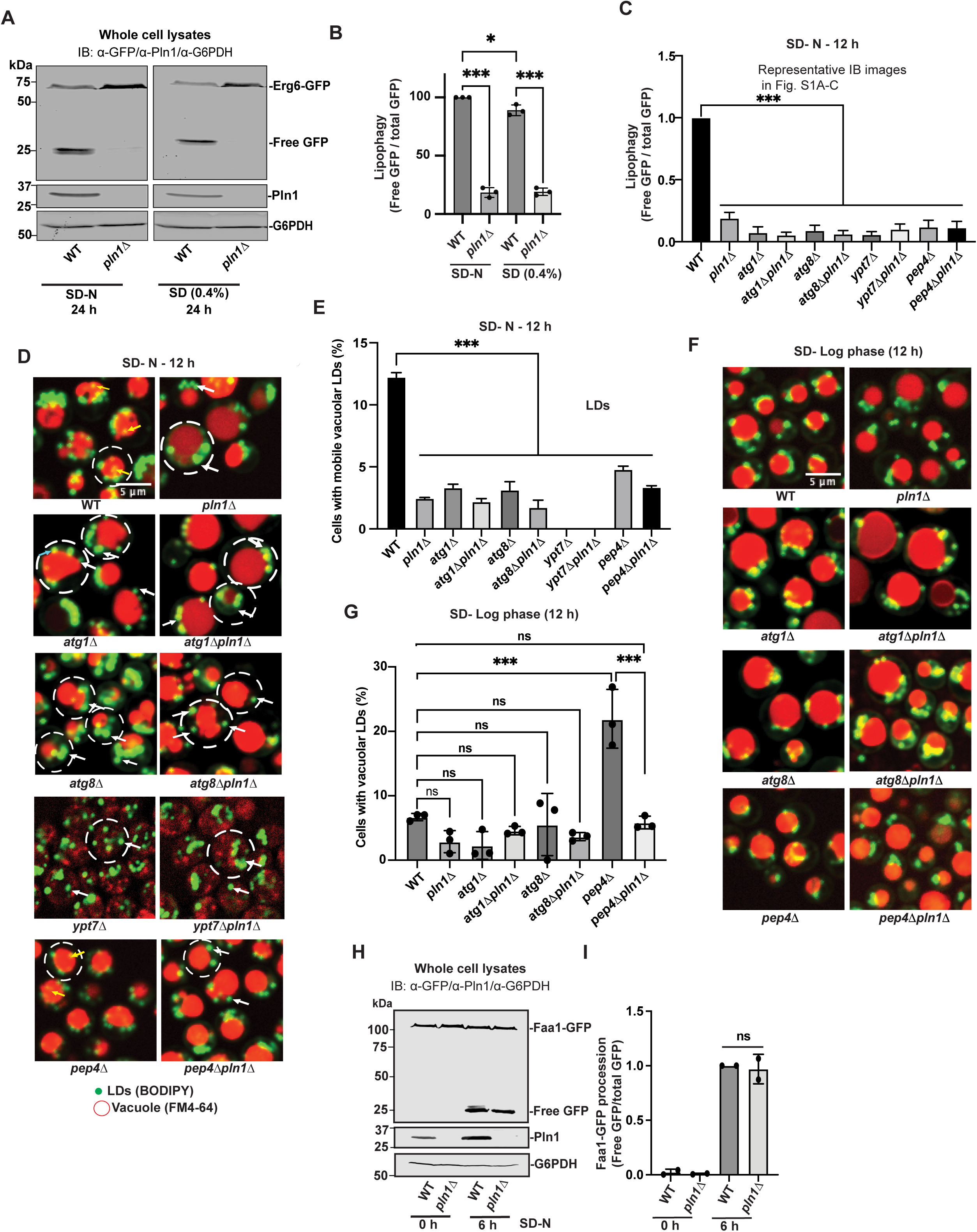
Pln1 is Essential for Lipophagy but Not General Autophagy. (**A-B**) Immunoblot for total protein extracts from WT and *pln1Δ* strains endogenously expressing Erg6-GFP collected after 24 h of starvation in (SD-N) or SD(0.4%) medium. (**A**) Representative images and (**B**) quantitative analysis. Bar graphs represent means ± SD (n = 3). **(C)** Immunoblot analysis of total protein extracts from WT, *pln1Δ, atg8Δ, atg8Δ pln1Δ, ypt7Δ, ypt7Δ pln1Δ, pep4Δ,* and *pep4Δ pln1Δ* strains endogenously expressing Erg6-GFP after 12 h of nitrogen starvation medium (SD-N). Representative images are in Fig. S1A-C. (**D**) Micrographs of indicated knockouts strains starved for 12 h in nitrogen free medium (SD-N) were stained with BODIPY for LDs and FM4-64 for vacuoles. Circles, cells; white arrow: cytosolic LDs; yellow arrow: LDs invaginated by vacuoles. (**E)** Quantification graph representing percentage of cells with mobile vacuolar LDs. (**F**) Micrographs of indicated knockouts strains grown up to log phase (12 h in SD medium). (**G**) Graph showing percentage of cells with vacuolar LDs. Bar graphs represent means ± SD (n = 3). **(H-I)** Immunoblots of total protein extracts from corresponding strains endogenously expressing Faa1-GFP after 6 h of nitrogen starvation (SD-N), **(H)** Representative image of immunoblot, **(I)** quantitative analysis. Bar graphs represent means ± SD (n = 3). ns, no significance; *, p ≤ 0.05; ***, p ≤ 0.001 (Ordinary one-way ANOVA, Tukey’s multiple comparisons test). Scale bar, 5 µm.

Next, we examined vacuolar LDs using confocal live-cell imaging. Vacuolar LDs are generally more mobile than cytosolic LDs, due to the loss of cytosolic interactions and their relatively small size — it has been suggested that lipophagy preferentially targets small LDs (Kang et al., 2024; Schott et al., 2019). Using this feature, we analyzed LDs (BODIPY-stained) in vacuoles (FM4-64 stained). These LDs appeared to move during Z-stack image acquisition (100 ms interval), in both wild-type and deletion strains. As expected, the presence of vacuolar mobile LDs was inhibited in cells harboring *atg1*Δ, *atg8*Δ, *pep4*Δ, or *ypt7*Δ (Fig. 1D-E). Importantly, the lipophagy defect in *pln1*Δ cells was as severe as in these strains, showing no obvious synthetic effects (Fig. 1C, 1E, N-starvation). During early stationary phase, *pep4*Δ promoted vacuolar accumulation of LDs presumably due to the role of Pep4 in precursor maturation of vacuolar proteinases (Fig. 1F-G). This accumulation in *pep4*Δ was prevented by *pln1*Δ. Supported by the Erg6-GFP reporter assay and cell imaging of vacuolar LDs, we concluded that Pln1 is a core regulator of lipophagy.

Nutrient starvation stimulates both bulk (non-selective) and selective autophagy, such as lipophagy and ER-phagy, the autophagy of the endoplasmic reticulum (ER) (Mochida et al., 2015; Ranganathan et al., 2022; Schuck et al., 2014), which collectively consumes Atg8 in the vacuole. We next utilized the 2×GFP-Atg8 processing assay to monitor total autophagy flux in wild-type and *pln1*Δ cells. As an LD-resident protein, we predicted that Pln1 would specifically affect autophagy directed toward LDs, with minimal impact on total autophagy flux. Indeed, after 6 hours of starvation in SD-N medium, the *pln1*Δ strain exhibited only a slight reduction in 2×GFP-Atg8 processing compared to wild-type cells (Fig. S1D-E). Moreover, ER-phagy, monitored by vacuolar processing of Faa1-GFP (an ER-localized protein), remained unchanged in *pln1*Δ cells (Fig. 1H-I). Hence, Pln1 specifically regulates lipophagy with negligible influence on general autophagy flux.

### Pln1 Ubiquitously Supports LD-Vacuole Association

Microlipophagy is the primary form of lipophagy in yeast, stimulated by glucose and nitrogen starvation (Seo et al., 2017; Zhang et al., 2020). Unlike macrolipophagy, where LDs are enclosed by autophagosomes, microlipophagy requires LD targeting, docking and tethering on the vacuole surface, which precedes the vacuolar invagination of LDs (Fig. 2A, illustration). Next, we examined at which step(s) Pln1 regulates microlipophagy.

**Figure 2.**
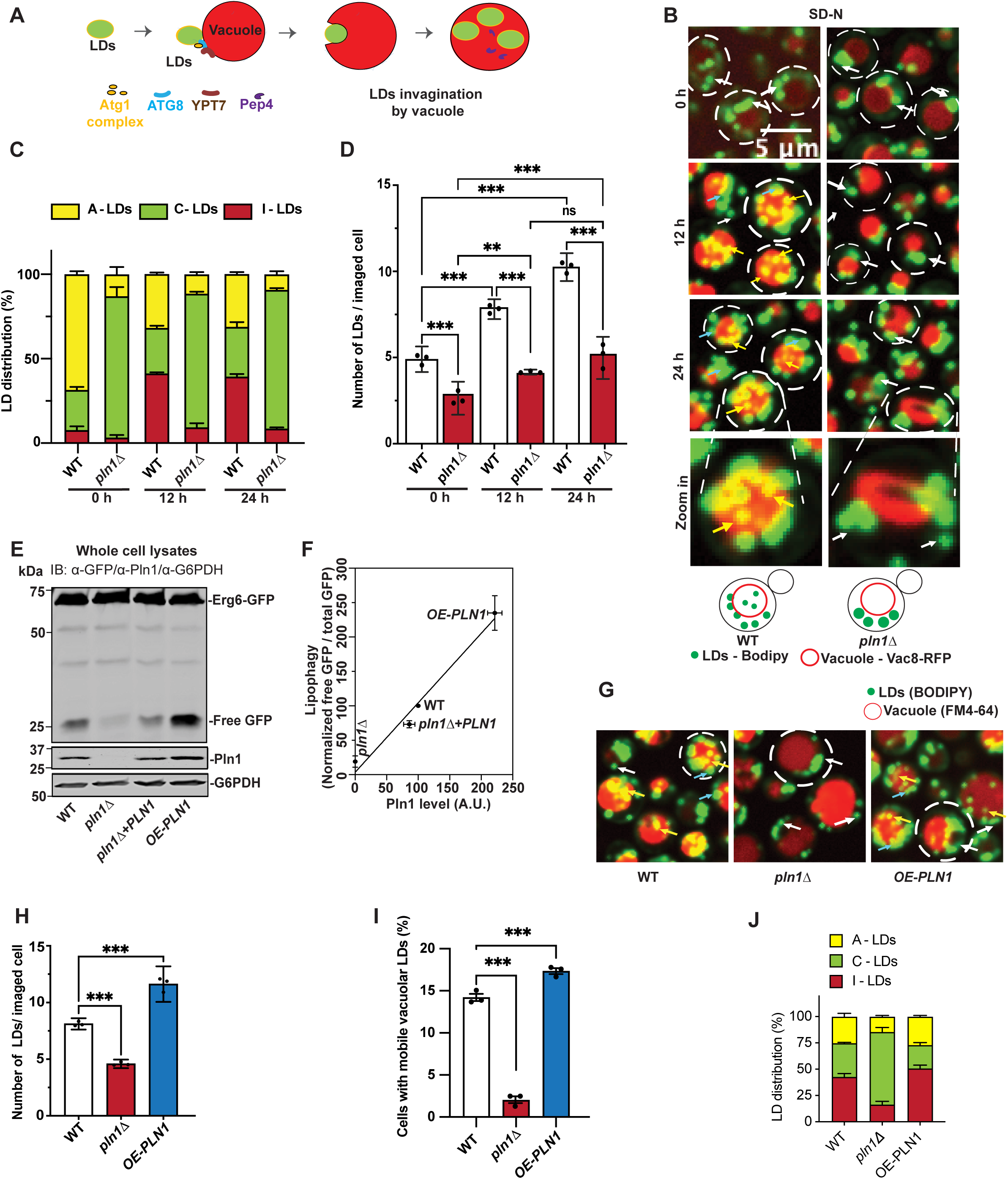
Pln1 Ubiquitously Supports LD-Vacuole Association. (**A**) Schematic of microlipophagy with regulators genetically deleted in this study. (**B**) Micrographs of cells with indicated genetic backgrounds analyzed at 12 h of SD-N starvation, stained with BODIPY for LDs and Vac8-RFP for vacuoles. (**C-D**) Quantification of (**B**), showing intracellular distribution of LDs, percentage of LDs that are invaginated by vacuoles (I - LDS), cytosolic, (C - LDs) or associated with vacuolar membrane (A-LDs) (**C**). Number of LDs imaged for cell (**D**). (**E**) Immunoblot of Erg6-GFP procession from whole cell lysates of WT, *pln1Δ* and other indicated strains; cells were collected after at 24 h of starvation in (SD-N) medium, **(F)** quantitative analysis plotting free GFP/ total GFP to Pln1 expression, normalized to wild-type (100). **(G)** Micrographs of cells with indicated genetic backgrounds analyzed at 12 h of SD-N starvation, stained with BODIPY for LDs and FM4=64 for vacuoles. Quantification of **(G)**, **(H)** number of LDs per cell **(I)** Percentage of cells with mobile vacuolar LDs. **(J)** showing intracellular distribution of LDs, percentage of LDs that are invaginated by vacuoles (I - LDS), cytosolic, (C - LDs) or associated with vacuolar membrane (A-LDs). Scale bar, 5 μm Bar graphs represent means ± SD (n = 3). **, p ≤ 0.01; ***, p ≤ 0.001 (Ordinary one-way ANOVA, Tukey’s multiple comparisons test). **(B, G)** Circles, cells; white arrow: cytosolic LDs; yellow arrow: LDs invaginated by vacuoles; blue arrow: LDs associated with vacuolar. Scale bar, 5 µm.

Using a spinning disk confocal fluorescent microscopy (FM) live-cell imaging approach, we investigated LD distribution in cells. BODIPY was used to stain LD lipids, and FM4-64 (or Vac8-RFP) was used for vacuoles. We classified LDs into three categories: intravacuolar LDs (I-LDs), vacuolar membrane-associated LDs (A-LDs), and cytosolic LDs (C-LDs), based solely on their location regardless of mobility. Due to the resolution of confocal cell imaging (∼250 nm), it is worth noting that A-LDs in our analysis include LDs tethered or partially invaginated in the vacuoles.

In wild-type cells, most LDs (∼70%) associate with vacuoles during late log phase (Fig. 2B-C, 0 h of strvation). Upon starvation (SD-N for 12 hours), there was a significant increase in the total number of LDs (Fig. 2D), and the proportion of I-LDs rose from ∼5% to ∼45%, accompanied by a decrease in A-LDs from ∼70% to ∼25%. This is consistent with A-LDs being an intermediate stage of microlipophagy. Remarkably, deletion of PLN1 (*pln1*Δ) greatly increased the proportion of C-LDs during late log phase and under both starvation conditions (Fig. 2B-C and Fig. S2A-E: WT: ∼30%, *pln1*Δ: ∼70-90% of total LDs), suppressing the cellular levels of A-LDs and I-LDs. In contrast, overexpression of Pln1 (OE-PLN1) enhanced lipophagy (Fig. 2E-F, Erg6-GFP processing; Fig. 2G-I, microscopy); and LD-vacuole association (Fig. 2J). Thus, Pln1 regulates microlipophagy as early as at the stage of LD-vacuole association.

Recent studies have shown that LD-resident proteins Ldo16 and Ldo45 bind the vacuolar membrane protein Vac8 to form an LD-vacuole tether during microlipophagy triggered by glucose starvation, though their role under nitrogen starvation remains unclear (Álvarez-Guerra et al., 2024; Diep et al., 2024). We found that deletion of LDOs (*ldo16/45*Δ) also led to the reduction of A-LDs under glucose starvation (13%), albeit to a lesser extent than *pln1*Δ (15%). The additional deletion of PLN1 further reduced LD-vacuole association (A-LDs) in *ldo16/45*Δ cells, but not vice versa (Fig. 3A-C). In addition, overexpression of Pln1 (OE-PLN1) did not rescue lipophagy or LD-vacuole association in *ldo16/45*Δ cells (Fig. 3A-C). Hence, productive LD-vacuole association during glucose starvation-triggered microlipophagy depends on both Pln1 and LDOs.

**Figure 3.**
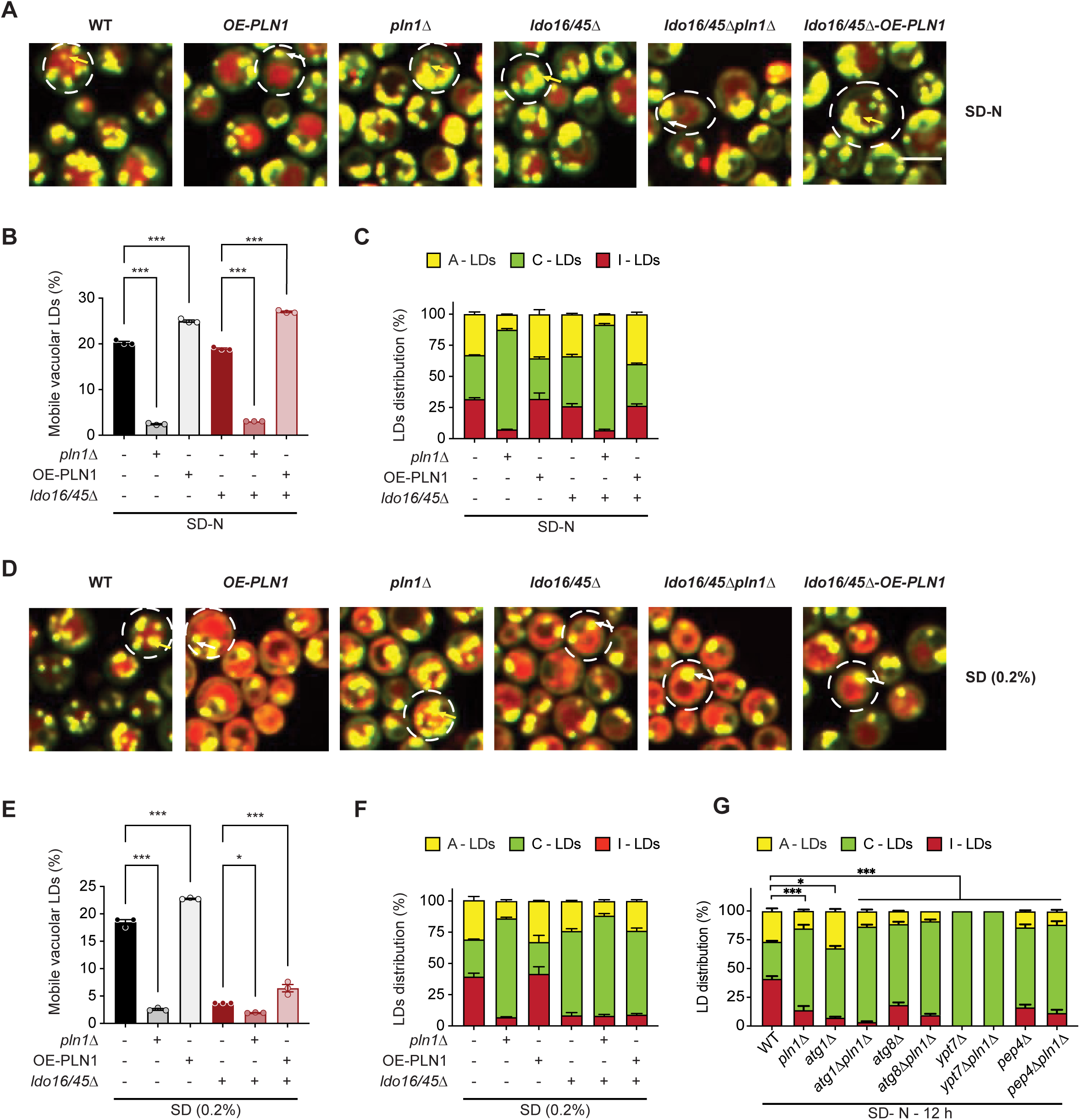
Identification of Pln1 Domains Essential for Its Role in Microlipophagy. **(A)** Micrographs of cells indicated knockouts strains grown in nitrogen starvation (SD-N) conditions were stained with BODIPY for LDs and FM4-64 for vacuoles. (**B-C)** Quantification graph representing **(B)** Percentage of cells with vacuolar LDs in corresponding stains and **(C)** percentage of intracellular distribution of LDs invaginated vacuoles-LDs (I - LDS), cytosolic, (C - LDs) and vacuolar membrane associated LDs, (A-LDs) in corresponding strains. **(D)** Micrographs of cells indicated knockouts strains grown in glucose starvation (SD [0.2%]) conditions were stained with BODIPY for LDs and FM4-64 for vacuoles. **(E-F)** Quantification graph representing **(E)** Percentage of cells with vacuolar LDs in corresponding stains and **(F)** percentage of intracellular distribution of LDs invaginated vacuoles-LDs (I - LDS), cytosolic, (C - LDs) and vacuolar membrane associated LDs, (A-LDs) in corresponding strains. **(G)** percentage of intracellular distribution of LDs in corresponding strains. percentage of LDs that are invaginated by vacuoles (I - LDS), cytosolic, (C - LDs) or associated with vacuolar membrane (A-LDs). Bar graphs represent means ± SD (n = 3). *, p ≤ 0.05; ***, p ≤ 0.001 (Ordinary one-way ANOVA, Tukey’s multiple comparisons test). **(A, D)** Circles, cells; white arrow: cytosolic LDs; yellow arrow: LDs invaginated by vacuoles. Scale bar, 5 µm.

Strikingly, unlike Pln1, LDO16/45 proteins are dispensable for microlipophagy under nitrogen starvation conditions (Fig. 3D-F). Thus, the LD-vacuole association factors involved in microlipophagy appear to be context-dependent: Pln1 is a ubiquitous regulator, whereas LDO16/45 specifically support tethering during glucose starvation. Hereafter, we focused the subsequent analyses primarily on nitrogen starvation-induced microlipophagy, unless stated otherwise, as the mechanism of LD-vacuole tethering under this condition remains poorly understood.

Atg1 and Atg8 support microlipophagy in yeast (Nakatogawa et al., 2009), while the underlying mechanism remains unclear. Intriguingly, we found that, similar to *pln1*Δ cells, *atg8*Δ cells exhibited reduced A-LDs under nitrogen starvation, whereas *atg1*Δ cells maintained wild-type levels of LD-vacuole association under nitrogen starvation (Fig. 3G). This observation suggests that Atg8, which is known to associate with vacuoles (Ishii et al., 2019; Munzel et al., 2021), might facilitate LD-vacuole association. Conversely, Atg1 may function downstream of this association step, potentially promoting vacuolar invagination of LDs.

Since lipophagy depends on vacuolar functionality, we next investigated whether a functional vacuole is essential for LD-vacuole association. Deletion of Pep4 (*pep4*Δ), a vacuolar proteinase required for post-translational precursor maturation of vacuolar proteinases, impairs vacuolar function (Ammerer et al., 1986; Jones et al., 1982) without altering vacuolar morphology. In *pep4*Δ cells, the proportion of A-LDs decreased to levels comparable to *pln1*Δ cells (Fig. 3G). Additionally, in *ypt7*Δ cells, characterized by fragmented vacuoles, measurable LD-vacuole association was essentially abolished (Fig. 3G). Consistently, Pln1 overexpression (OE-PLN1) was unable to rescue either lipophagy or A-LD populations in *atg1*Δ, *ypt7*Δ, or *pep4*Δ cells (Fig. S2F-I). These findings collectively indicate that LD-vacuole association during microlipophagy strictly requires autophagy machinery and a functional vacuole.

### Identification of Pln1 Domains Essential for Its Role in Microlipophagy

The Pln1 protein contains a characteristic PAT domain (residues P_16_ to V_92_), which is found in most perilipins (PLNs) (Bickel et al., 2009; Gao et al., 2017). The PAT domain is named after the three founding members of this protein family: perilipin (PLN1), Adipophilin (PLN2), and TIP47 (PLIN3) (Londos et al., 2005; Yamaguchi, 2007). Starting from the N-terminus, the PAT domain of Pln1 is followed by a hydrophobic region (residues G_112_ to L_131_) and a Repeat domain (residues Y_190_ to N_226_), which is flanked by two gap regions GAP1 (residues R_132_ to N_189_) and GAP2 (residues E_227_ to E_263_ ) (Fig. 4A) (Gao et al., 2017). To identify the domains and sequences critical for Pln1’s role in lipophagy, we generated strains expressing truncated Pln1 variants (Fig. 4A), driven by either the endogenous *PLN1* promoter or the *PGK1* promoter (OE) (for enhanced expression), all at the PLN1 genomic locus.

**Figure 4.**
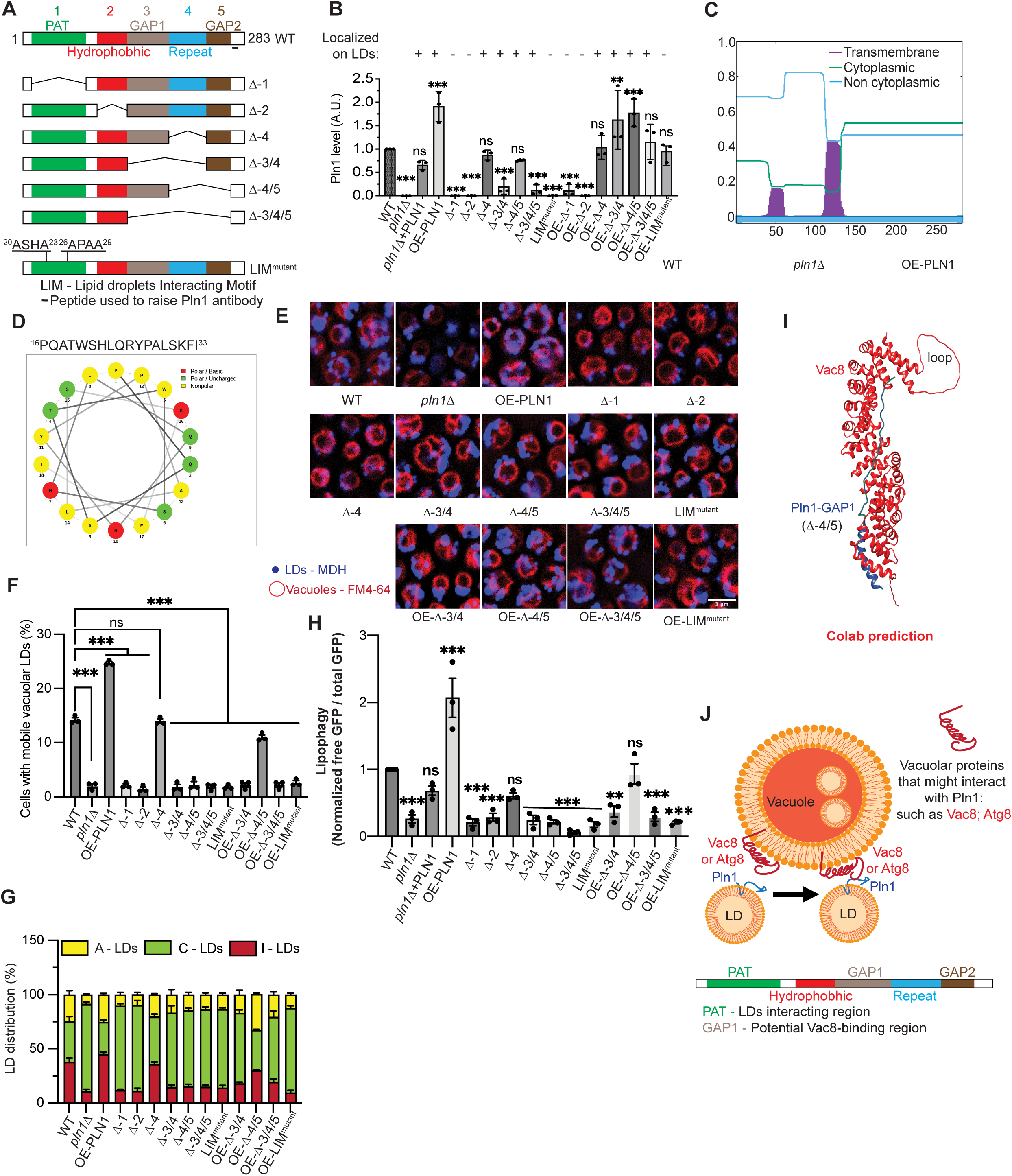
Pln1 Supports Microlipophagy of LDs regardless of Neutral Lipid Composition. (**A**) Schematic representation of PLN1 variants examined in this study. (**B**) Quantification of protein levels of indicated Pln1 variants. Data from Fig. S3A-B, S4A-B. The symbols above the bars are analysis on co-localization of Pln1 variants (C-terminal tagged by mCherry) with BODIPY-stained LDs. Data from Fig. S3C-D. +: co-localized; -: no signal or not co-localized. **(C)** Hydrophobicity plot of Pln1 protein sequence using Phobius. **(D)** a helix prediction by Netwheels of putative sequences of 18 aa in length including regions of LIM1 and LIM1 motifs in PAT domain of Pln1p. (**E-G**) Fluorescent microscopy of indicated Pln1 variants analyzed at 12 h of SD-N starvation. Shown are (**E**) micrographs stained with MDH for LDs and FM4-64 for vacuoles (Scale bar, 3 µm), quantitative analysis of (**F**) percentage of cells with mobile vacuolar LDs and (**G**) percentage of distribution of intracellular LDs invaginated vacuoles-LDs (I - LDS), cytosolic, (C - LDs) and vacuolar membrane associated LDs, (A-LDs).in corresponding strains. (**H**) Quantification of lipophagy marked by Erg6-GFP procession in indicated strains. Data from Fig. S3A-C. Bar graphs represent means ± SD (n = 3). (**I**) Predicted interaction between Vac8 and *Pln1-GAP1*, modeled by ColabFold. (**J**) Model of Pln1-mediated LD-vacuole association. PAT domain of Pln1 facilitates Pln1 to reside on LDs, whereas GAP1 domain of Pln1 might interact with Vac8 that resides on vacuoles. ns, no significance; **, p ≤ 0.01; ***, P ≤ 0.001 (Ordinary One-way ANOVA, Tukey’s multiple comparisons test).

Among the mutants, the Δ*PAT (*Δ*-1)* and Δ*Hydrophobic (*Δ*-2)* proteins were almost undetectable by immunoblotting, even under the *PGK1* promoter, while reduced protein levels of Δ*GAP1/*Δ*Repeat (*Δ*-3/4)* and Δ*GAP1/*Δ*Repeat/*Δ*GAP2 (*Δ*-3/4/5)* could be restored to wild-type levels or above with the *PGK1* promoter (OE) (Fig. 4B, Fig. S3A-B). When detectable, the Pln1 truncation mutants mostly resided on LDs, suggesting a potential link between Pln1’s stability and its localization to LDs (Fig. 4B, Fig.S3C-D). Therefore, we hypothesize that the PAT and Hydrophobic domains of Pln1 mediate its localization to LDs, which in turn stabilizes Pln1.

The PAT domain in mammalian PLNs is crucial for targeting and anchoring PLNs to the surface of LDs (Orlicky et al., 2008; Rowe et al., 2016). Our sequence analysis of Pln1 identified an ∼18 amino acid motif (16-PQATWSHLQRYPALSKFI-33) within the PAT domain with mixed hydrophobic and positively charged residues, which could form a cationic amphipathic helix capable of interacting with the phospholipid monolayer of LDs (Fig. 4C-D). Strikingly, mutating four hydrophobic residues in this motif (W_20_xxL_23_ and Y_26_xxL_29_) to alanine (Pln1-LIM^mutant^) completely abolished localization of Pln1 to LDs ( Fig. 4B; S3C-D) and stimulated its degradation (Fig. 4B; S3A). While the *PGK1* promoter increases Pln1-LIM^mutant^ protein levels to levels close to endogenous Pln1 (Fig.4B; S3B), Pln1-LIM^mutant^ (OE) remained in the cytosol, failing to localize to LDs (Fig. 4B; S3D). We conclude that the PAT domain of Pln1 contains an LD-interacting motif (LIM) that is essential for its LD localization.

Remarkably, Pln1-LIM^mutant^ (OE) cells were as defective as *pln1*Δ in LD-vacuole association and lipophagy, as shown by cell imaging (Fig. 4E-G) and Erg6-GFP processing (Fig. 4H; S4A). This suggests that LD localization is a prerequisite for Pln1’s role in these processes. On the other hand, Δ*-3/4* (OE), Δ*-4/5* (OE), and Δ*-3/4/5* (OE) strains retained LD localization (Fig. 4B, Fig. S3D), but only Δ*-4/5* (OE) cells exhibited levels of LD-vacuole association and lipophagy close to wild-type (Fig. 4F; S4C). By process of elimination, these results support the notion that the GAP_1_ region (Fig. 4A, #3; residues R_132_ to N_189_) regulates LD-vacuole association. As GAP_1_ is dispensable for Pln1’s LD localization, we conclude from these data that LD residency alone is necessary but not sufficient for Pln1 to promote LD-vacuole association.

Based on AlphaFold2 prediction, GAP1 consists of an intrinsically disordered region (IDR, residues 132-165), followed by an α-helix (residues 166-189). Intriguingly, a recent study revealed that the IDR regions of Ldo16 and Ldo45, two LD proteins, interact with the vacuolar surface protein Vac8 to promote LD-vacuole contact site formation (Álvarez-Guerra et al., 2024). We hypothesize that part of the GAP_1_ IDR might bind Vac8 to assist LD-vacuole association, either through or independent of LDOs. Using ColabFold, we found that all five top models of Vac8-GAP_1_ interaction showed part of Pln1’s GAP_1_ IDR (residues 136-165) in contact with the minor groove of the Vac8 superhelix, the same groove that binds LDO proteins (Álvarez-Guerra et al., 2024) (Fig. 4I). Given the apparent competition between GAP_1_ and LDOs for Vac8 binding, one scenario is that Pln1 and LDOs bind to Vac8 sequentially to support distinct steps of LD-vacuole association. Alternatively, LD-vacuole association might involve Pln1, LDOs, Vac8 and other factors, e.g., Atg14 and Atg6 (Lei et al., 2021), in a higher-order complex to support high-affinity LD-vacuole interaction.

Overall, our data support the following model (Fig. 4J): The N-terminus of Pln1 – the PAT domain and a hydrophobic segment – is essential for ensuring its residency on LDs. This region likely anchors it to the surface of LDs by direct interaction with lipids (Fig. 4C, TM prediction). Toward the C-terminus, GAP_1_ facilitates LD-vacuole association, possibly involving interactions with vacuolar surface proteins such as Vac8. In the simplest-case scenario, Pln1 functions as part of a tether, pairing with Vac8; although, other possibilities exist. This hypothesis and its alternatives will be discussed further later.

### Pln1 Supports Microlipophagy of LDs regardless of Neutral Lipid Composition

Lipid droplets (LDs) within cells vary in size and lipid composition, giving rise to distinct subpopulations (Cartwright et al., 2015; Gao et al., 2017; Speer et al., 2023). In addition to triacylglycerol (TAG), LD cores contain steryl esters (SE), and are surrounded by a phospholipid monolayer coated with diverse LD-associated proteins (Sorger et al., 2004). TAG biosynthesis is catalyzed primarily by the enzymes Dga1 and Lro1 (Oelkers et al., 2002; Oelkers et al., 2000), whereas SE biosynthesis involves Are1 and Are2 (Zweytick et al., 2000). Consequently, genetic manipulation of these enzymes results in LDs enriched specifically in TAG or SE. For example, the triple mutant strains *lro1*Δ *are1*Δ *are2*Δ (3KO-DGA1) and *dga1*Δ *lro1*Δ *are2*Δ (3KO-ARE1) predominantly produce LDs composed of TAG and SE, respectively (Fig. 5A, left illustration) (Sandager et al., 2002).

**Figure 5.**
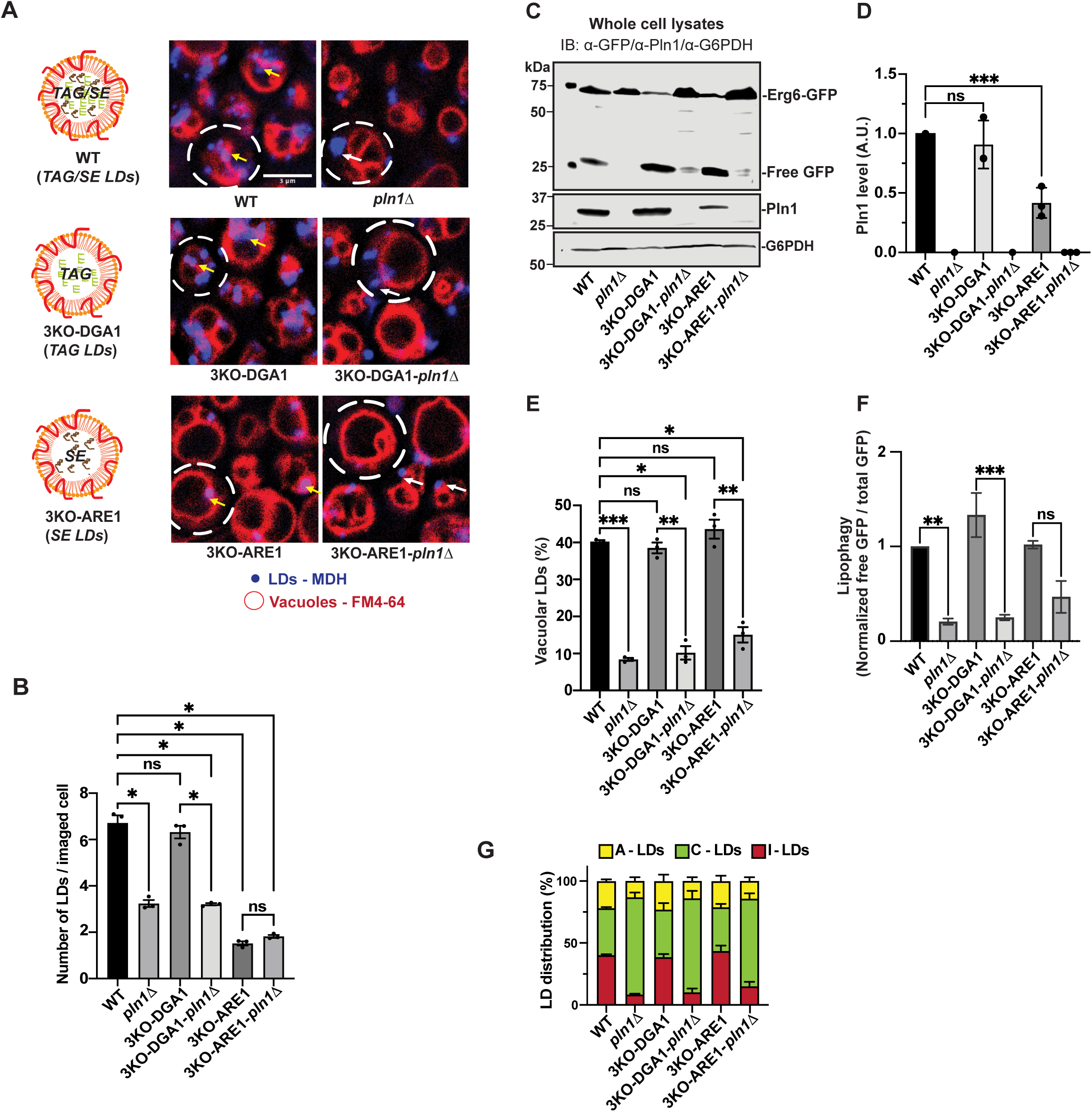
Pln1-Assisted TAG Biogenesis, Not Lipophagy, Determines Intracellular TAG Levels During Nitrogen Starvation. **(A)** Schematic representation of LDs with various types of neutral lipids (left side): WT – LDs with both TAG and SE, 3KO-DGA1-LDs with TAG and 3KO-ARE – LDs with SE; **right,** confocal micrographs for WT, *pln1Δ,* 3KO-DGA1, 3KO-DGA1-*pln1Δ,* 3KO-ARE1, and 3KO-ARE1-*pln1Δ* strains, starved under nitrogen starvation medium (SD-N) for 12 h; cells were stained by MDH for LDs and FM4-64 for vacuoles. Circles, cells; white arrow: cytosolic LDs; yellow arrow: LDs invaginated by vacuoles. Scale bar, 3 μm. **(B)** Analysis of data from (**A**). Graph representing number of LDs per imaged cell. **(C-D)** Immunoblot analysis of whole cell lysates from WT, *pln1Δ,* 3KO-DGA1, 3KO-DGA1-*pln1Δ,* 3KO-ARE1, and 3KO-ARE1-*pln1Δ* strains endogenously expressing Erg6-GFP; cells were collected after 12 h of starvation in (SD-N) medium. Shown are (**C**) immunoblot images, (**D**) quantitative analysis of Pln1 levels. Membranes were probed with antibodies directed against GFP, Pln1 and G6PDH; G6PDH is used as loading control. **(E)** Analysis of data from **(A)**, percentage of vacuolar LDs in indicated strains. (**F**) Quantification of Erg6-GFP processing in indicated strains in immunoblot shown in **(C)**. **(G)** percentage of distribution of intracellular LDs invaginated vacuoles-LDs (I - LDS), cytosolic, (C - LDs) and vacuolar membrane associated LDs, (A-LDs) in corresponding strains. Bar graphs represent means ± SD (n=3). ns, no significance; *, p ≤ 0.05; **, p ≤ 0.01; ***, p ≤ 0.001 (Ordinary one-way ANOVA, Tukey’s multiple comparisons test).

Previous work estimated that Pln1 occupies approximately 15–86% of the LD surface (Gao et al., 2017). Moreover, Pln1 preferentially associates with TAG-rich LDs, although it also binds to SE-rich LDs at lower but detectable levels (Gao et al., 2017). These observations prompted us to investigate whether Pln1 predominantly regulates lipophagy of TAG-rich LDs. Using fluorescence imaging, we found similar numbers of LDs in wild-type (WT) and 3KO-DGA1 (TAG-rich LD) cells, whereas LD abundance was significantly lower in 3KO-ARE1 (SE-rich LD) cells (Fig. 5A-B), which also exhibited a ∼50% reduction in intracellular Pln1 levels (Fig. 5C-D). Despite differences in LD abundance, lipophagy occurred robustly in WT, 3KO-DGA1, and 3KO-ARE1 strains, and—surprisingly—was strictly dependent on Pln1, as determined by cell imaging (Fig.5E) and Erg6-GFP processing assays (Fig. 5F). Moreover, deletion of *PLN1* (*pln1*Δ) similarly reduced the proportion of A-LDs in WT, 3KO-DGA1, and 3KO-ARE1 strains (Fig.5G). These results indicate that even reduced levels of Pln1 associated with SE-rich LDs are sufficient to mediate microlipophagy. Thus, the neutral lipid composition (TAG vs. SE) does not significantly influence the Pln1-dependent regulation of lipophagy.

### Pln1-Assisted TAG Biogenesis, Not Lipophagy, Determines Intracellular TAG Levels During Nitrogen Starvation

Pln1 plays dual supportive roles in LD metabolism, facilitating both LD biogenesis (through enhancing TAG synthesis) (Gao et al., 2017). and LD degradation via microlipophagy. To understand how these roles influence intracellular TAG levels during nutrient starvation, we measured TAG abundance in wild-type and *pln1*Δ cells under nitrogen-starvation conditions (SD-N) for up to 36 hours (Fig. 6A). To distinguish Pln1’s role in TAG synthesis from its function in lipophagy, we included *atg1*Δ cells, which block lipophagy but not TAG biogenesis. Besides lipophagy, cytosolic lipolysis— primarily mediated by Tgl3, Tgl4, and Tgl5 in yeast—also significantly contributes to TAG turnover (Athenstaedt & Daum, 2003, 2005; Klein et al., 2016). Therefore, we examined the effect of deleting these lipases (*tgl3/4/5*Δ) on TAG levels to assess their role in LD homeostasis during nitrogen starvation (Fig. 6A).

**Figure 6.**
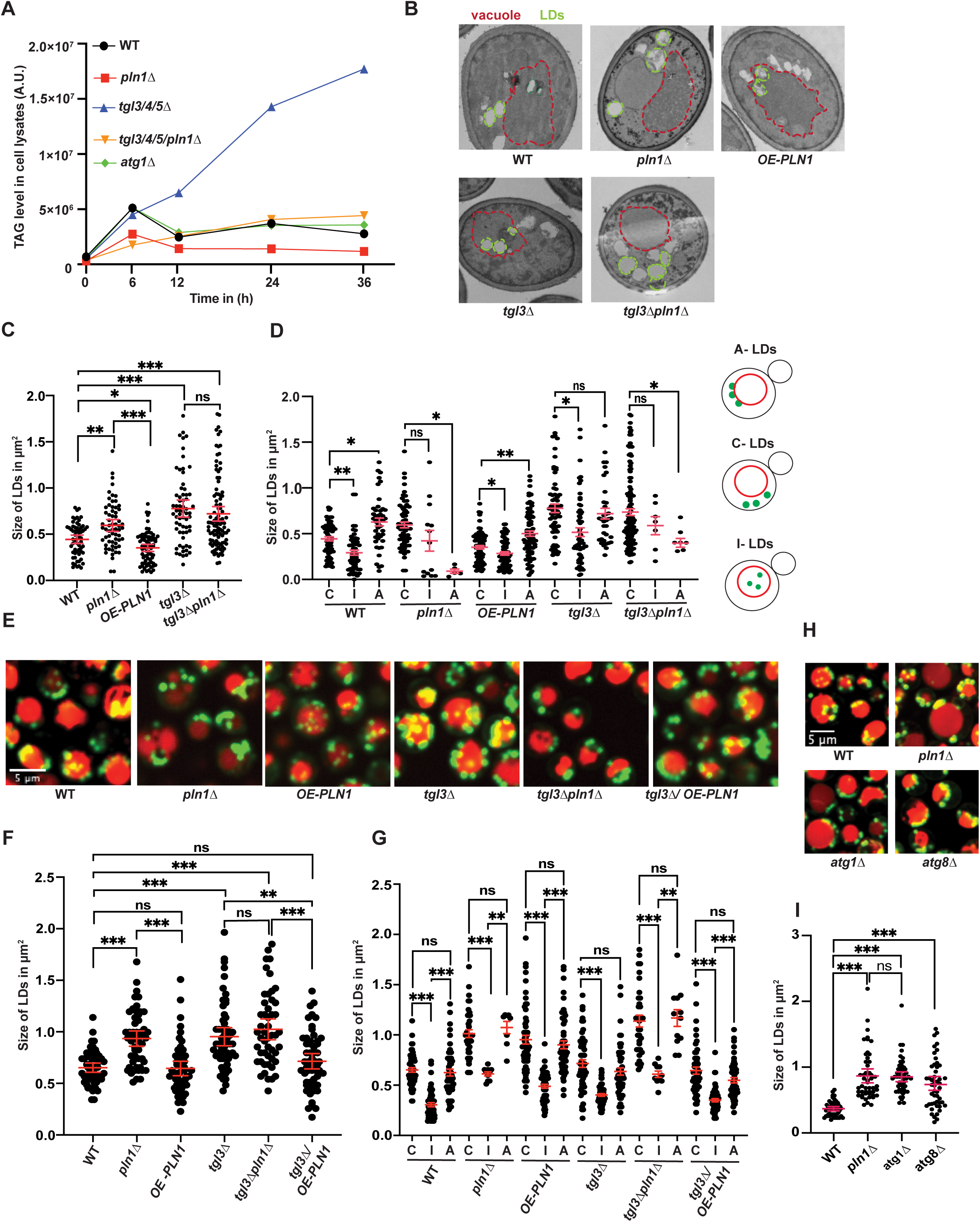
Pln1-Mediated Lipophagy Regulates Intracellular LD Size and Distribution. **(A)** Quantification of TAG content in whole cell lysates from WT, *pln1Δ, tgl3Δ tgl4Δ tgl5Δ (tgl3/4/5Δ*), *tgl3Δ tgl4Δ tgl5Δ pln1Δ (tgl3/4/5/ pln1Δ*), and *atg1Δ* strains starved in SD-N medium for different time intervals (0 h, 6h, 12 h, 24h and 36 h), using HPLC. **(B)** Electron microscopy (EM) images of cells with indicated genetic backgrounds, after 12 h of SD-N starvation. **(C)** Quantification of (**B**), showing sizes of total LDs (left) and **(D)** Showing the size of LDs after sorting LDs, based on interaction vacuole (right). **(E)** Micrographs of WT and various other indicated mutant strains after 12 h of SD-N starvation, LDs were stained with BODIPY and FM4-64 for vacuoles. **(F)** Quantification of size of LDs in corresponding stains of (**E**) and **(G)** quantification of size of LDs in vacuoles (I – LDs), in cytosol (C – LDs) and in association with vacuolar membrane (A – LDs) in corresponding strains from confocal imaging. (**H**) Micrographs of WT, *pln1Δ, atg1Δ* and *atg8Δ* strains after 12 h of SD-N starvation, LDs were stained with BODIPY and FM4-64 for vacuoles. (**I**) Quantification of (**H**), showing sizes of LDs in indicated stains. Scale bar, 5 μm. Quantification of size of LDs in corresponding stains. (Each dot represents a LD (n = 50). ns, no significance; *, p ≤ 0.05; **, p ≤ 0.01***, p ≤ 0.001 (Brown-Forsythe and Welch ANOVA tests, Dunnett’s T3 multiple comparisons test). Scale bar, 5 µm.

Previous studies reported that nitrogen starvation initially stimulates LD biogenesis, storing TAG and SE, with LD formation peaking after approximately 6 hours (Li et al., 2015). Subsequently, prolonged starvation induces TAG and SE consumption through lipolysis and lipophagy, promoting LD turnover to sustain cell survival. As expected, intracellular TAG levels were initially low during late log phase (Fig. 6A, 0 h of starvation), with *pln1*Δ cells exhibiting the lowest levels. After 6 hours of nitrogen starvation, TAG content increased in all strains, albeit to a lesser extent in *pln1*Δ cells, indicating ongoing TAG and LD biogenesis (Fig. 6A). Importantly, at this early stage, blocking TAG turnover through lipophagy (*atg1*Δ) or lipolysis (*tgl3/4/5*Δ) did not further increase TAG accumulation beyond wild-type levels. Additionally, deletion of lipases (*tgl3/4/5*Δ) did not elevate TAG levels in *pln1*Δ cells, suggesting minimal TAG turnover at this initial stage.

Between 12 and 36 hours of nitrogen starvation, TAG accumulated significantly in the *tgl3/4/5*Δ cells, indicative of active cytosolic lipolysis. Surprisingly, despite active lipophagy observed during the same period (Fig.1A-B;1D-E), inhibition of lipophagy by *atg1*Δ or *pln1*Δ did not lead to intracellular TAG accumulation, in sharp contrast to *tgl3/4/5*Δ (Fig. 6A). This result indicates that cytosolic lipolysis, rather than lipophagy, is primarily responsible for TAG degradation under these conditions. Notably, the substantial TAG accumulation observed in *tgl3/4/5*Δ cells was reversed by *pln1*Δ (Fig. 6A). Thus, it is Pln1-assisted TAG biogenesis—not lipophagy—that predominantly determines intracellular TAG levels during nitrogen starvation.

### Pln1-Mediated Lipophagy Regulates Intracellular LD Size and Distribution

The minimal impact of lipophagy inhibition on intracellular TAG levels was unexpected, especially given clear evidence that LDs were delivered to vacuoles (Fig. 1D-E), and Erg6-GFP and Pln1 were degraded between 12 and 36 hours of starvation (Fig. 1A-B; Fig. S5A-B). Nonetheless, consistent with our observations, previous studies have reported a similarly mild effect of lipophagy on TAG content during nutrient starvation (Álvarez-Guerra et al., 2024).

Given its minimal impact on total TAG content, we next investigated how lipophagy influences intracellular LDs. In the absence of lipophagy (in *pln1*Δ or *atg8*Δ), the intracellular LD distribution shifted towards cytosolic accumulation (C-LDs), reducing the share of A-LDs and I-LDs (Fig. 3G). In addition, we observed that LD size significantly increased in *pln1*Δ cells, sharply contrasting with cells overexpressing Pln1 (OE-PLN1), as revealed by electron microscopy (EM) (Fig. 6B-D) and fluorescence microscopy (Fig. 6E-G). Remarkably, deletion of *PLN1* enlarged intracellular LDs to sizes comparable to those observed upon the loss of Tgl3-mediated cytosolic lipolysis. This similarity was particularly striking since *tgl3/4/5*Δ cells accumulate intracellular TAG, whereas *pln1*Δ cells exhibit dramatically reduced TAG levels (Fig. 6A). These results led us to hypothesize that Pln1-mediated lipophagy, similar to lipolysis, is involved in controlling LD size.

To test this hypothesis, we analyzed LD size in *atg1*Δ cells. In contrast to *pln1*Δ, deletion of *ATG1* blocks lipophagy (Fig. 1C; 1E) without affecting TAG biosynthesis or homeostasis (Fig. 6A). Remarkably, we found that LDs in *atg1*Δ (and *atg8*Δ) cells were similarly enlarged, closely resembling those observed in *pln1*Δ cells (Fig.6H-I), suggesting that blocking LD autophagy was the cause. Mechanistically, we noted that vacuolar LDs (I-LDs) were generally smaller than cytosolic LDs (C-LDs and A-LDs) (Fig. 6D; 6G), consistent with previous reports indicating that lipophagy preferentially delivers smaller LDs into the vacuole (Kang et al., 2024; Schott et al., 2019). We speculate that lipophagy regulates LD size by consuming a portion of a large LD, as is the case for liver (Schulze et al., 2020), and/or selectively transferring small LDs to vacuoles, thereby limiting their growth into larger LDs in the cytosol. Based on these findings, we conclude that lipophagy can regulate the size and intracellular distribution of LDs, without leading to TAG accumulation.

### Pln1 Regulates Long-Term Quiescent Cell Survival

Upon nutrient depletion, cells transition from proliferation to quiescence to prioritize survival. The successful survival of quiescent cells relies on the coordinated recycling of cytoplasmic components for essential building blocks (e.g., amino acids) and energy production (e.g., ATP). This coordination involves bulk autophagy (Green et al., 2011; Yen & Klionsky, 2008) and lipid consumption (Hardie & Carling, 1997; Zechner et al., 2012). Given the special role of Pln1 in both TAG synthesis and lipophagy, we investigated its function in quiescent cell survival under nitrogen starvation and acute glucose starvation (SD, 0.2% glucose).

Consistent with previous reports (Sturgeon et al., 2019), autophagy-deficient *atg1*Δ or *atg8*Δ cells exhibited reduced survival after approximately nine days of nitrogen starvation compared to wild-type cells (Fig. 7A), confirming the essential role of autophagy in recycling cytoplasmic components. However, the specific contribution of lipophagy remained unclear. Interestingly, *pln1*Δ cells showed no detectable survival disadvantage under these conditions, whereas overexpression of Pln1 (*pPGK1-PLN1*) slightly reduced cell survival compared to wild-type (Fig. 7A; S6A-C). Similar trends were observed during acute glucose starvation, although cells generally survived longer (∼two weeks longer) and the loss of *ATG1* or *ATG8* had only mild effects on survival (Fig. 7A). Thus, lipophagy appears not critical for long-term quiescent cell survival.

**Figure 7.**
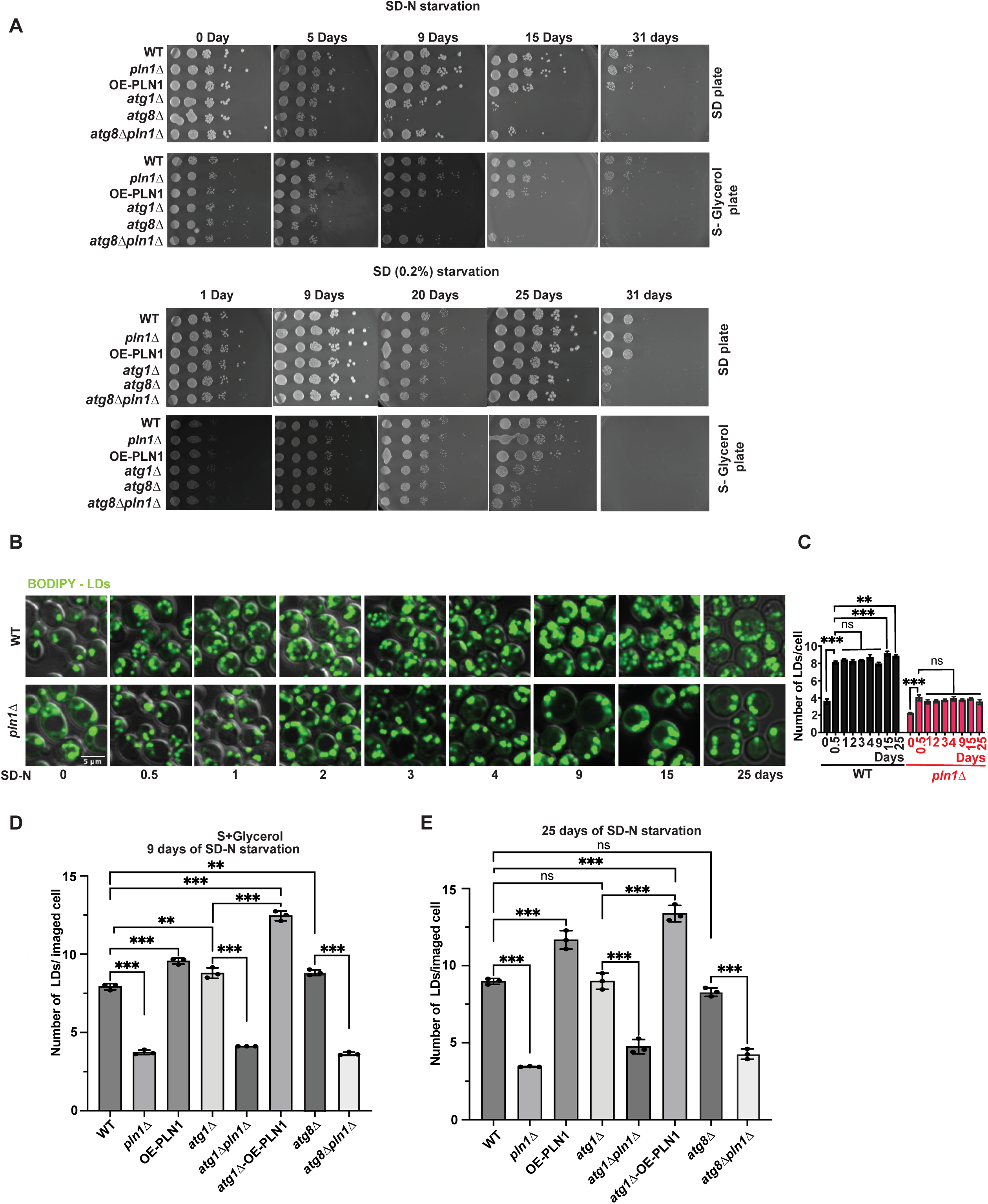
Pln1 Regulates Long-Term Quiescent Cell Survival. **(A)** Survival studies of WT, *pln1Δ, OE-PLN1, atg1Δ, atg8Δ, atg8Δ pln1Δ* strains under different time intervals of starvation from 0 days to 31 days in (top) SD-N or (bottom) SD (0.2%) media, and survival studies were monitored by spotting on different media condition: SD medium and Glycerol medium. **(B)** Micrographs of WT and *pln1Δ* strains in SD-N starvation medium, LDs were stained with BODIPY for different time intervals (0 h, 0.5, 1 day, 2 days, 3 days, 4 days, 9 days, 15 days and 25 days). **(C)** Quantification of (B) showing LDs number per imaged cell. **(D-E)** Quantification of LDs number in WT, *pln1Δ, OE-PLN1, atg1Δ, atg1Δpln1Δ, atg1Δ-OE-PLN1, atg8Δ* and *atg8Δpln1Δ* strains starved for 9 days **(D)** and 25 days **(E)** in SD-N starvation medium. Bar graphs represent means ± SD (n = 3). ns, no significance; *, p ≤ 0.05; **, p ≤ 0.01; ***, p ≤ 0.001; (Ordinary one-way ANOVA, Tukey’s multiple comparisons test). Scale bar, 5 µm.

Nevertheless, one caveat is that deletion of *PLN1* reduces cellular LDs and TAG to approximately 50% of wild-type levels due to impaired TAG synthesis (Fig. 6A, TAG level; Fig.2D, LD counting). Intriguingly, in both wild-type and *pln1*Δ cells, the intracellular LD numbers remained stable even after 25 days of nitrogen starvation (Fig. 7B-C). Consequently, we speculated that lipophagy might primarily facilitate survival only when intracellular LD and TAG levels are at wild-type levels, and that reduced intracellular TAG and LD content might bypass the requirement for lipophagy for cell survival. To test this hypothesis, we generated double mutants (*pln1*Δ *atg8*Δ and *pln1*Δ *atg1*Δ) to lower LD biogenesis (*pln1*Δ) in autophagy-defective backgrounds (*atg8*Δ and *atg1*Δ). Supporting our hypothesis, deletion of *PLN1* further decreased LD content in *atg8*Δ cells (Fig. 7D, 9 days in SD-N; Fig. 7E, 25 days in SD-N) and partially rescued their survival defect (Fig. 7A).

The *pln1*Δ mediated partial rescue in autophagy-defective (*atg8*Δ) cells is context dependent. Notably, *atg1*Δ had a stronger negative effect on cell survival compared to *atg8*Δ in liquid medium (Fig. S6A-C), possibly due to autophagy-independent Atg1 kinase activities. This may explain why *pln1*Δ did not enhance survival in *atg1*Δ cells (Fig. 7A; S6A-C). Another interesting finding in our study is that *pln1*Δ enhanced survival in *atg8*Δ cells only under nitrogen starvation, not under acute glucose starvation (Fig. 7A). It was suggested that autophagy is essential for survival during nitrogen starvation, while being dispensible during glucose starvation (Adachi et al., 2017), in line with our finding that *atg8*Δ *or atg1*Δ *moderately* reduces cell survival after glucose restriction. Therefore, although the mechanism remains unknown, it seems that Pln1 regulates cell survival in the context of activation of autophagy specifically programmed by nitrogen starvation (see discussion).

Collectively, our findings demonstrate a ubiquitous role of Pln1 in regulating lipophagy at the stage of LD-vacuole association, under various conditions. We show evidience that Pln1-regulated LD homeostasis coordinates with bulk autophagy, regulating quienscent cell viability under nitrogen starvation.

## Discussion

### Mechanism of Pln1-Dependent LD-Vacuole Association

Pln1 plays a dual role in LD biology, supporting both LD biogenesis (Gao et al., 2017) and microlipophagy, as demonstrated in this study. In mammals, five perilipin proteins (PLINs 1–5) critically regulate LD dynamics, with PLIN3 being the closest homolog to yeast Pln1. Although the PAT domain is a defining feature of perilipins, the intrinsically disordered region (IDR) of Pln1, essential for LD-vacuole association and lipophagy, does not share sequence similarity with other mammalian PLIN proteins. While mammalian PLIN2 and PLIN3 can partially restore TAG accumulation in yeast *pln1*Δ strains (Gao et al., 2017; Hammoudeh et al., 2023), their Pln1-like role in supporting microlipophagy in mammalian cells remains to be determined. Moreover, whereas PLIN2 and PLIN3 protect LDs from lipase activity (Bickel et al., 2009; Cinato et al., 2024; Londos et al., 2005), it is uncertain whether yeast Pln1 possesses a similar protective function. Thus, it remains unclear if the role of Pln1 in macrolipophagy and LD turnover is conserved.

How Pln1 facilitates LD-vacuole association is not fully understood. The vacuolar surface protein Vac8 binds Ldo16 and Ldo45, promoting LD-vacuole tethering (Álvarez-Guerra et al., 2024). Interestingly, Ldo16 and Ldo45 bind a specific structural groove on Vac8, predicted in our study to also interact with Pln1’s IDR. Additionally, Pln1’s localization to LDs is indispensable for LD-vacuole association and microlipophagy, suggesting Pln1 may cooperate with Ldo16 and Ldo45 to stabilize LD-vacuole tethering complexes. Notably, while Ldo proteins are dispensable during nitrogen starvation, Pln1 supports microlipophagy under diverse conditions, implying the existence of distinct tethering mechanisms responsive to different nutritional cues, all sharing Pln1 as a central regulator.

Pln1 may additionally interact with Ypt7, the yeast homolog of mammalian Rab7 GTPase, essential for vacuole integrity, functionality, and autophagy (Haas et al., 1995; Shvarev et al., 2022; Wada et al., 1992; Wichmann et al., 1992; Wickner & Rizo, 2017). Proteomic studies identified potential Pln1-Ypt7 interactions via co-immunoprecipitation with Ypt7 as bait, analyzed by LC-MS/MS (Bouchez et al., 2015). Thus, Pln1 might engage with Ypt7 to facilitate LD docking on vacuoles, analogous to Ypt7’s interaction with Vps39 during mitochondria-vacuole interactions in vCLAMP formation (Acoba et al., 2020; González Montoro et al., 2018).

Alternatively, Pln1 may regulate LD-vacuole associations indirectly. Pln1 is highly abundant on LDs (Gao et al., 2017), and our data indicate that elevated Pln1 levels during starvation negatively impact the LD association of proteins such as Erg6. Thus, modulating Pln1 abundance could alter the LD proteome in important ways, influencing the spatial distribution of LD proteins including Ldo16 and Ldo45. Pln1 likely plays multiple roles at the LD-vacuole interface, dynamically interacting with various factors such as LDOs and Atg14 to fine-tune tethering events.

Importantly, we identified Pln1 as a crucial regulator of lipophagy under nitrogen starvation (SD-N) and glucose restriction—conditions known to activate distinct signaling pathways. However, lipophagy can also be induced under various other stress conditions, including stationary phase or ER stress induced by DTT or tunicamycin (Garcia et al., 2021; Kang et al., 2024; van Zutphen et al., 2014; Zhang et al., 2020). The roles of nutrient-sensing pathways such as mTOR and AMPK in Pln1-mediated lipophagy remain open questions that warrant further investigation.

### The Physiological Role of Lipophagy

Vacuoles (lysosomes) are enriched with hydrolytic enzymes capable of degrading biological polymers such as proteins, lipids, nucleic acids, and polysaccharides (Bonam et al., 2019; De Duve et al., 1955; Wang et al., 2018). It has long been assumed that lipophagy leads to the vacuolar (lysosomal) degradation of triacylglycerol (TAG), the primary lipid component of lipid droplets (LDs). However, known yeast TAG lipases, such as Tgl2, Tgl3, Tgl4, and Tgl5, specifically target cytosolic LDs (Athenstaedt & Daum, 2003, 2005; Jandrositz et al., 2005; Klein et al., 2016; Kurat et al., 2006), and no vacuolar TAG lipase activity has been reported. The only characterized lipase in yeast vacuoles, Atg15, primarily functions as a phospholipase (Kagohashi et al., 2023; Watanabe et al., 2023). Thus, future studies are required to conclude whether vacuolar lipases for neutral lipids, including TAG and SE exist in yeast.

Our study revealed that intracellular TAG levels remain relatively stable in wild-type (WT) cells during prolonged nitrogen starvation (Fig. 6A). Concurrently, LDs are delivered to vacuoles, resulting in the degradation of LD-resident proteins such as Erg6 and Pln1 (Fig. 1). Consistent with stable TAG levels, LD numbers per cell remain relatively unchanged, even after approximately 25 days of nitrogen starvation (Fig. 2D, 7D-E). One possible explanation is that active LD and TAG biogenesis compensates for TAG loss via lipophagy or lipolysis. Inhibiting lipophagy in *pln1*Δ or *atg1*Δ cells did not result in TAG accumulation compared to WT. In contrast, TAG accumulated significantly in lipolysis-defective *tgl3*Δ*/4*Δ*/5*Δ cells (Fig. 6A), suggesting lipophagy may not be the primary regulator of intracellular TAG homeostasis under these conditions.

Interestingly, defects in lipophagy led to increased LD size during starvation. Yeast LDs typically range from approximately 200 to 800 nm in diameter in WT cells (W303-1A strain). Previous studies indicated smaller LDs are more susceptible to lipophagy, possibly due to their higher propensity for autophagosome engulfment or vacuolar membrane indentation and rupture (Ivanovska et al., 2023; Schott et al., 2019; van Zutphen et al., 2014). Consistent with these reports, we observed smaller LDs within vacuoles compared to cytosolic or vacuolar membraneassociated LDs, though LD-vacuole association itself showed no size preference (Fig. 6D, 6G). We hypothesize that lipophagy controls cytosolic LD size by selectively transferring smaller LDs into vacuoles, thus preventing their fusion into larger droplets in the cytosol. We also cannot rule out that the yeast vauole may consume a portion of a large LD, as has been shown in macrolipophagy in liver (Schulze et al., 2020). Although the physiological significance of LD size regulation during starvation remains uncertain, altered LD size correlates with changes in intracellular LD distribution (Fig. 6D, 6G), potentially affecting their stability, composition, and overall functionality (Kang et al., 2024).

A fundamental question in quiescent cell survival is how bulk autophagy and lipid consumption pathways coordinate. Autophagy recycles cellular components, whereas lipid consumption provides energy. Remarkably, deletion of *PLN1* (*pln1*Δ) enhanced survival in nitrogen-starved *atg8*Δ cells. Comparing *pln1*Δ *atg8*Δ with *atg8*Δ cells, LD content was significantly reduced in the double deletion due to impaired LD biogenesis, while autophagy (including lipophagy) remained absent in both cases. Thus, lower LD content appears beneficial for survival under nitrogen starvation, especially when autophagy is compromised. Mechanistically, we speculate that lipid consumption via lipolysis or lipophagy (for energy production) needs to be coordinated with bulk autophagy (for amino acids and other building blocks) so that essential synthesis activities in the quienscient cells can be maintained. Alternatively, digestion of neutral lipids generate fatty acids and other toxic lipid metabolites, potentially causing cellular damage that can be mitigated by autophagic recycling. Thus, the reduced survival in *atg8*Δ cells may result from impaired recycling coupled with increased accumulation of harmful lipid metabolites. Consequently, the loss of Pln1 reduces levels of energy production, as well as toxic lipid intermediates by lowering LD and TAG content, thereby improving cell viability under autophagy-deficient conditions.

### Limitations

We acknowledge that it is technically chanllenging to fully discriminate I-LDs from A-LDs, due to the resolution of the spinning disk confocal microscopy ( ∼250 nm).

## Materials and Methods

### Yeast strains and genetics

All yeast strains used in this study are based on W303-1A (MATa leu2-3/112 ura3-1 trp1-1 his3-11/15 ade2-1 can1-100) (Thomas and Rothstein, 1989). All yeast strains used in this study are listed in Tables S1. All yeast deletion strains and yeast strains with endogenous tagging of genes with GFP or mCherry are genetically modified using standard homologous recombination methods and transformation methods. Oligonucleotides used in this study are listed in Table S2. The plasmids encoding Pln1 were created by PCR amplification of respective fragments and assembling them by Gibson assembling method with overlapping PCR and resulting plasmid sequences were listed (Folder - PLN1 and perilipin-containing plasmids). The final constructs were introduced into the pBMF4 plasmid backbone using HindIII restriction site followed by 661 base pairs of 5’PLN1-UTR followed by PLN1 coding fragment along with ADH1 terminator and TEF promoter and terminator containing HIS3 along with 3’UTR of PLN1 with SacII restriction enzymes and transformed into respective strains for integration. PLN1△-1, PLN1△-2, PLN1△-4, PLN1△-3and 4, PLN1△-4 and 5, PLN1△-3,4 and 5, LIM1 mutant and LIM2 mutant and LIM mutants plasmids were constructed in the same way by amplifying the corresponding fragments from the pJMG4-PLN1 plasmid and fusing them by overlapping PCR, preserving the promoter, To create the LIM1 variants carrying point mutations in the putative LDs Interacting Motifs, the point mutations were introduced in respective oligonucleotides, and the amplified fragments containing the mutated sites were joined with overlapping PCR. The genomic single deletion of *PLN1* (*pln1*△) was generated, briefly the open reading frame coding for Pln1 was first substituted with a cassette containing hygromycin cassette using *pFA6-hygMX6 CYC1* term as a template.

### Yeast culturing conditions

All strains used in this study were precultured overnight in minimal synthetic dextrose (SD) medium (0.67% Yeast Nitrogen Base with ammonium sulfate, 0.2% histidine, 0.2% uracil, 0.4% adenine, 0.12% lysine, 0.6% leucine, and 0.4% tryptophan) containing 2% glucose, at 30°C by shaking at 200 rpm. For nitrogen starvation conditions, overnight cultures from single colony were back diluted with 0.02 OD_600_ of cells in SD medium and grown up to 1.0 OD_600_ for 12 to16 h, and cells are washed with three times with distilled water and resuspended in SD-N medium with 3 OD_600_/ml for required time intervals. For Acute Glucose Restriction (AGR) conditions, overnight cultures in SD were washed and reinoculated into SD (0.4% glucose), with 3 OD_600_/ml for indicated time intervals. For deletion and tagging of genes, yeast cells were grown in rich medium (YPD) containing 20 g/l peptone (Gibco Bacto BD Bio-sciences), 10 g/l yeast extract (Bacto BD Biosciences) and 2% glucose. For subsequent selection of mutants, YPD plates containing hygromycin B (HYG5000), G418 (Sigma-Aldrich, A1720-5G) and for deletion and tagging of genes using auxotrophic selection, SD plates with all amino acids except for histidine, or tryptophan, or leucine, were used.

### Protein extraction for immunoblotting

30 OD_600_ of cells were collected by centrifugation and resuspended in 10% trichloroacetic acid, for overnight at -80° C and next day cells were collected by centrifugation and whole-cell extracts for immunoblots were prepared by resuspending cell pellet in 2× Laemmli sample buffer (Laemmli, 1970) and neutralizing with 10 μl to 15 μl of 5N NaOH, followed by vertexing with acid-washed glass beads on the MiniBeadBeater 16 (model 607; Biospec Products) for three 1-min pulses. Samples were boiled for 10 min and then centrifuged for the same duration at 13,000 rpm to remove insoluble material and glass beads. The supernatant fraction was collected in fresh tube and the identical cell equivalents (typically 1.0 OD_600_ units) were subjected to SDS-PAGE and western blotting. Immunoblotting was performed by probing membranes with antibodies against the GFP-epitope (dilution 1:5000, mouse, Roche life sciences 1181446001), anti-G6PDH (dilution 1:10000, rabbit, Sigma, A9521), and in house generated anti-PLN1 antibody (dilution 1:5000, rabbit) (Gao, Q et al. 2017), as well as respective LiCOR-fluorescent-conjugated secondary antibodies against mouse (dilution 1:10000, mouse, IRDye^®^ 680RD Goat anti-Mouse IgG Secondary Antibody 926-68070) or rabbit (dilution 1:10000, rabbit, LICOR IRDye^®^ 680RD Goat anti-Rabbit IgG Secondary Antibody 926-68071). LICOR Odyssey detection system was used for detection. Densitometric quantification of immunoblots was performed with Image Studio lite software.

### Quantification of total cellular triacylglycerol levels by using HPLC

10 OD_600_ of cells were harvested, washed with distilled water and cell pellet was resuspended in 300 μl of distilled water with glass beads and lysed with a MiniBeadBeater 16 (model 607; Biospec Products) using 3 cycles of 1 min pulse and 1 min rest. Then 800 μl of distilled water was added to the tube and vortexed for resuspending followed by transfer of 800 μl cell lysate to conical shape screw cap glass tube (Kimble® 73785-10) and another round of washing with 800 μl of distilled water was performed and added to previous collection. 4 ml of methanol and 2 ml of chloroform was added to the total cell lysate collected and vortexed for 1 minute. Followed by addition of 2 ml of 0.9% NaCl and vortex for 1 minute. Top aqueous layer was removed with paster pipette after centrifugation at 1500 rpm for 2 minutes at room temperature. Then additional wash was given to organic layer with 2 ml of 2M KCl followed by 1 minute of vertexing and centrifugation at 1500 rpm for 2 minutes. Remove the top aqueous layer and transfer the bottom organic layer to new glass tube and dry under liquid nitrogen. Total lipids were resuspended in 100 μl of hexane and lipids were resolved for 15 minutes on Agilent ZORBAX CN-5 μm HPLC column with inner diameter of 4.6 mm and 150 mm column length. Using solvent systems of from Hexane: Tert-Butyl Methyl Ether (TBME) (97:3) and from 10 minutes onwards Hexane: TBME (80:20) solvent system. Concentration of triacylglycerol was measures and quantified using standards TAG concentrations curves.

### Analysis of cellular survival during nitrogen starvation and analysis of cell growth

Starvation induced cell death was measured by analyzing rate of cell survival. Cultures from respective starvation days were collected and washed and diluted to 0.1 OD of cells with 1 ml of SD medium. Using 250 µl of 0.1 OD (10^-1^) as starting culture and serially diluted up to 10^-2^, 10^-3^, 10^-4^, and 10^-5^ in 96-well microplates with clear, flat bottom (Greiner Bio-ONE). Plates were shaking at 200 rpm at 28 C and growth was measured by monitoring OD600 every 15 minutes for 3 days using a plate reader (2300 EnSpire, Perkin Elmer). This experiment is designed to track how the cells respond to starvation over several days, with the key observation being the rate at which cells survive (reflected by OD_600_). The survival rate is indicated by how well cells grow after serial dilution, where reduced growth in higher dilutions suggests decreased viability due to starvation-induced cell death.

### Fluorescence Microscopy

1 OD_600_ of cells is harvested at designated time points. The cells are stained using specific dyes to visualize certain cellular components. BODIPY^493/503^ or MDH (Monodansylpentane) is used for staining lipid droplets (LDs) and FM 4-64: is used for vacuolar staining. 1 μL of BODIPY^493/503^ stock (*1* mg/mL) *dissolved* in DMSO and *1 µl of* FM^TM^ 4-64 stock of *(1* mg/mL) *dissolved* in DMSO *were* added directly to *1 ml of* culture media and Incubate*d* in the dark on a rocker for 30 minutes. Cells are centrifuged, and the supernatant is removed. Cells are resuspended in 100 µl of water or 1xPBS. And finally, 7 ul of resuspended cells were used for imaging. The microscope hardware and image acquisition methods, including projections from z-stacks, were as previously described using Zeiss LSM700 (Gao et al., 2017), with the modification that the z-stack consisted of 25 images spaced 0.35 μm apart. Image acquisition was performed using Slidebook.6.0.4 (Intelligent Imaging Innovations). Alternatively, cells were imaged on a Zeiss confocal microscope, the system included 483-, and 515-nm laser lines, and Slidebook software.6.0.4 (Intelligent Imaging Innovations).

### Image analysis and quantification

Images obtained with the ZEISS LSM700 were first processed using the Slidebook 6.0.4 software. The open-source software ImageJ was used to further process and quantify all confocal micrographs. To process the images, mid plains were selected and Images from the same experiment were processed with similar settings. Brightness and contrast were adjusted for each channel equally in individual experiments for analysis. The distribution of number of LDs present in the cell were classified based on association with the vacuole. Initially percentage of number of cells showing vacuolar LDs were calculated to total number of cells. Lipophagy was calculated manually measuring number of LDs ‘inside of vacuoles and outside of the vacuole and LDs associated with vacuolar membranes. ‘Relative percentage of lipophagy’ was obtained by calculating the I-LDs in respective strains. LD size was quantified manually by measuring the area of LDs occupied using ImageJ. Number of LDs per cell was quantified by manually counting of the segmented LDs.

### Transmission electron microscopy

For transmission electron microscopy, samples were prepared as described (Wright 2000). Briefly, yeast strains were collected after 24 hours of starvation and washed with 1xPBS, cells were fixed with 4% paraformaldehyde and serially washed with 10% ethanol, 20% ethanol, 30% ethanol, 40% ethanol, 50% ethanol, 60% ethanol, 70% ethanol, 80% ethanol, 90% ethanol, 95% ethanol and 100% ethanol respectively. Cells were resuspended with 100% ethanol. After embedding, samples of 70–90-nm thickness were placed on 200 mesh copper Formvar grids and poststained. The thin sections were observed in a transmission electron microscope (1400 plus; JEOL) AMT BIOSPRINT16M-ActiveVu mid mount CCD camera, Serial EM tomography and montaging software. Size of the lipid droplets were quantified manually by measuring the area of LDs occupied using ImageJ.

### Structural modelling

To model the conservation of the Vac8 surface, a multiple sequence alignment was made that focussed on Vac8 only using 2 rounds of HHblits, which searches into a nr30 database (non-redundant above 30% identity). After the first iteration 106 hits were included with e-value <10^-20^. After the second iteration, 294 proteins were included (e-value < 10^-42^). This excluded other Armadillo repeat proteins. Aligned sequences were submitted along with the solved structure of Vac8 to the ConSurf server to obtain conservation scores for residues on the surface scaled between 0-10. To model the interaction of Vac8 with Ldo16, ColabFold was seeded with Vac8 residues (560 residues, 19-578, missing the disordered N-terminus) and full length Ldo16 (148 residues). The structure shown in Figure 4I is the rank 1 model, for which a version was obtained with side-chains positioned in relaxed conformations.

### Statistical analyses

Data are presented as bar graphs used to represent data where the height of the bars corresponds to the mean values of the data in different experimental groups or conditions. Means ± SD (n = 3). The asterisks (***) indicate a very significant result, with a p-value of less than 0.001. One-Way ANOVA (Analysis of Variance) is used to compare means across multiple groups to determine if there are any statistically significant differences between them. Tukey’s Multiple Comparisons post hoc test applied after a one-way ANOVA to identify which specific groups differ from each other. Sample size, referring to independent biological replicates, is indicated in the respective figure legends. For quantification of size of LDs in corresponding stains. Each dot represents a LD (n = 50). ns, no significance; *, p ≤ 0.05; **, p ≤ 0.01; ***, P < 0.001; (Brown-Forsythe and Welch ANOVA tests, Dunnett’s T3 multiple comparisons test) is applied for quantification of size of LDs. All statistical analysis was performed using GraphPad Prism.

## Acknowledgement

The authors would like to thank Mike Henne for expertise and reagents that lead us to study lipid droplets in cell. We thank J. Friedman, M. Henne, and members of the Wang lab for comments on the manuscript.

We are grateful to Department of Cell Biology for providing microscopy facility. We thank UTSW Core Sequencing Facility and the UTSW Core Electron Microscopy Facility for their technical support.

This work was supported by a grant from the National Institutes of Health to F. Wang (R01GM133899), to Joel M. Goodman (GM084210), as well as funding from Nancy Cain and Jeffrey A. Marcus Scholar in Medical Research, in Honor of Dr. Bill S. Vowell, to F. Wang.

## Author Contribution Statement

Conceptualization, J.P., J.M.G, and F.W.; Methodology, J.P., A.T., W.P., J.M.G and F.W.; Investigation, J.P., B.F., J.R., A.T., J.L., J.M.G. and F.W.; Formal Analysis, J.P., R.Z. and F.W.; Visualization, J.P., R.Z. and F.W. Writing-Original Draft, J.P.; Writing-Review & Editing, J.P., J.M.G., W.P. and F.W.; Funding Acquisition, J.M.G. and F.W.

---

**Figure S1. Pln1 is required for efficient degradation of LD protein Erg6 but not general autophagy and ERphagy under nitrogen starvation.**

**(A-C)** Immunoblots of total protein extracts from corresponding strains endogenously expressing Erg6-GFP after 12 h of nitrogen starvation (SD-N) **(A)** WT, *pln1Δ, OE-PLN1, atg1Δ, atg1Δ pln1Δ, atg1Δ-OE-PLN1.* **(B)** WT, *pln1Δ, OE-PLN1, atg8Δ, atg8Δ pln1Δ, ypt7Δ, ypt7Δ pln1Δ, ypt7Δ-OE-PLN1,* **(C)** WT, *pln1Δ, OE-PLN1, pep4Δ, pep4Δ pln1Δ, pep4Δ-OE-PLN1, tgl3Δ, tgl3Δ pln1Δ, tgl3Δ-OE-PLN1* strains **(D-E)** Immunoblots of total protein extracts from corresponding strains endogenously expressing 2×GFP-Atg8 after 6 h of nitrogen starvation (SD-N), **(D)** Representative image of immunoblot, **(E)** quantitative analysis. Bar graph represents mean ± SD (n = 3). **, p ≤ 0.01 (Ordinary one-way ANOVA, Tukey’s multiple comparisons test).

**Figure S2. Pln1 is required for efficient lipophagy under both nitrogen starvation and glucose restriction.**

**(A)** Micrographs of strains with indicated genetic backgrounds analyzed at 12 h of SD-N starvation, and **(B)** SD (0.2%) medium from 0 h to 48 h and LDs were stained with BODIPY for LDs and FM4-64 for vacuoles. **(C)** percentage of LDs that are invaginated vacuoles-LDs (I - LDS), cytosolic, (C - LDs) and vacuolar membrane associated LDs, (A-LDs). **(D)** Graph representing percentage of cells with vacuolar LDs in WT and *pln1Δ* strains grown in (SD-N) and SD (0.2%) medium from 0 h to 48 h. **(E)** Graph representing percentage of cells with mobile vacuolar LDs in WT and *pln1Δ* strains grown in (SD-N) and SD (0.2%) medium from 0 h to 48 h. **(F)** Micrographs of cells with indicated genetic backgrounds analyzed at 12 h of SD-N starvation, stained with BODIPY for LDs and FM4=64 for vacuoles. Quantification of **(F)**, **(G)** number of LDs per cell **(H)** Percentage of cells with mobile vacuolar LDs. **(I)** showing intracellular distribution of LDs, percentage of LDs that are invaginated by vacuoles (I - LDS), cytosolic, (C - LDs) or associated with vacuolar membrane (A-LDs). Scale bar, 5 μm. Bar graphs represent means ± SD (n = 3). *, p ≤ 0.01; ***, P ≤ 0.001; (Ordinary one-way ANOVA, Tukey’s multiple comparisons test). **(A, F)** Circle, cells; white arrow: cytosolic LDs; yellow arrow: LDs invaginated by vacuoles; blue arrow: LDs associated with vacuoles. Scale bar, 5 µm.

**Figure S3. Molecular dissection of Pln1p and Identification of Pln1 domains essential for LDs targeting and maintaining stability of Pln1**

**(A-B)** Immunoblot showing the stability and expression profile of Pln1 in WT and various Pln1 mutant strains with *PLN1-*promoter **(A)** or overexpression with *PGK1-*promoter **(B)**. **(C)** Localization studies of C-terminal tagging of Pln1 and various other Pln1 mutants. WT (wild type as negative control), Pln1-mCherry, Δ-4-mCherry, Δ-3/4-mCherry, Δ-4/5-mCherry -mCherry, Δ-3/4/5-mCherry -mCherry, and LIM^mutant^-mCherry under *PLN1-* promoter (*pPLN1*-) and *PGK1-* promoter (*pPGK1-*). Circles, cells. **(D)** Co-localization events of mCherry tagged Pln1 and other mutants with LDs stained with BODIPY. Gray value of the BODIPY (green) and mCherry (red) signals along the white lines in magnified images were analyzed.

**Figure S4. Identification of Pln1 Domains Essential for its Role in Lipophagy.**

Representative immunoblot showing Erg6-GFP procession in whole cell lysates; cells expressing Erg6-GFP with indicated genetic backgrounds were collected from 12 h of SD-N starvation.

**Figure S5. Pln1-Assisted degradation of LDs proteins.**

**(A)** Immunoblotting for Pln1-mCherry expression and stability for protein extracts from WT-Pln1-mCherry strain under SD-N starvation for various time interval’s (0, 6, 12, 24 and 48 h). **(B)** Graph indicating ratio of Pln1-mCherry to total mCherry indicating the stability of Pln1p. Bar graph represents mean ± SD (n = 3). ns, no significance; *, p ≤ 0.01 (Ordinary one-way ANOVA, Tukey’s multiple comparisons test).

**Figure S6. Growth curves of strains treated by N-starvation.**

Growth curves of WT, *pln1Δ, OE-PLN1, atg1Δ, atg1Δ pln1Δ, atg1Δ-OE-PLN1, atg8Δ* and *atg8Δ pln1Δ* strains after nitrogen starvation for indicated days **(A)** 0 days of starvation. **(B)** 15 days of starvation. and **(C)** 25 days of starvation.

**Table S1. *Saccharomyces cerevisiae* strans used in this study.**

All strains used in this study are derived from W303-1A background (MATa; *leu2-3,112*; *trp1-1*; *can1-100*; *ura3-1*; *ade2-1*; *his3-11,15*).

**Table S2. DNA oligos used in this study.**

Sequences are 5’ to 3’. KO, knock-out; Fwd, forward primer; Rev, reverse primer.

## References

Acoba, M. G., Senoo, N., & Claypool, S. M. (2020). Phospholipid ebb and flow makes mitochondria go. J Cell Biol, 219(8). 10.1083/jcb.202003131

Adachi, A., Koizumi, M., & Ohsumi, Y. (2017). Autophagy induction under carbon starvation conditions is negatively regulated by carbon catabolite repression. J Biol Chem, 292(48), 19905–19918. 10.1074/jbc.M117.817510

Álvarez-Guerra, I., Block, E., Broeskamp, F., Gabrijelčič, S., Infant, T., de Ory, A., Habernig, L., Andréasson, C., Levine, T. P., Höög, J. L., & Büttner, S. (2024). LDO proteins and Vac8 form a vacuole-lipid droplet contact site to enable starvation-induced lipophagy in yeast. Dev Cell, 59(6), 759–775.e755. 10.1016/j.devcel.2024.01.014

Ammerer, G., Hunter, C. P., Rothman, J. H., Saari, G. C., Valls, L. A., & Stevens, T. H. (1986). PEP4 gene of Saccharomyces cerevisiae encodes proteinase A, a vacuolar enzyme required for processing of vacuolar precursors. Mol Cell Biol, 6(7), 2490–2499. 10.1128/mcb.6.7.2490-2499.1986

Arlt, H., Sui, X., Folger, B., Adams, C., Chen, X., Remme, R., Hamprecht, F. A., DiMaio, F., Liao, M., Goodman, J. M., Farese, R. V., & Walther, T. C. (2022). Seipin forms a flexible cage at lipid droplet formation sites. Nature Structural & Molecular Biology, 29(3), 194–202. 10.1038/s41594-021-00718-y

Athenstaedt, K., & Daum, G. (2003). YMR313c/TGL3 encodes a novel triacylglycerol lipase located in lipid particles of Saccharomyces cerevisiae. J Biol Chem, 278(26), 23317–23323. 10.1074/jbc.M302577200

Athenstaedt, K., & Daum, G. (2005). Tgl4p and Tgl5p, two triacylglycerol lipases of the yeast Saccharomyces cerevisiae are localized to lipid particles. J Biol Chem, 280(45), 37301–37309. 10.1074/jbc.M507261200

Athenstaedt, K., & Daum, G. (2006). The life cycle of neutral lipids: synthesis, storage and degradation. Cellular and Molecular Life Sciences CMLS, 63(12), 1355–1369. 10.1007/s00018-006-6016-8

Bickel, P. E., Tansey, J. T., & Welte, M. A. (2009). PAT proteins, an ancient family of lipid droplet proteins that regulate cellular lipid stores. Biochim Biophys Acta, 1791(6), 419–440. 10.1016/j.bbalip.2009.04.002

Bonam, S. R., Wang, F., & Muller, S. (2019). Lysosomes as a therapeutic target. Nature Reviews Drug Discovery, 18(12), 923–948. 10.1038/s41573-019-0036-1

Bouchez, I., Pouteaux, M., Canonge, M., Genet, M., Chardot, T., Guillot, A., & Froissard, M. (2015). Regulation of lipid droplet dynamics in Saccharomyces cerevisiae depends on the Rab7-like Ypt7p, HOPS complex and V1-ATPase. Biol Open, 4(7), 764-775. 10.1242/bio.20148615

Cartwright, B. R., Binns, D. D., Hilton, C. L., Han, S., Gao, Q., & Goodman, J. M. (2015). Seipin performs dissectible functions in promoting lipid droplet biogenesis and regulating droplet morphology. Mol Biol Cell, 26(4), 726–739. 10.1091/mbc.E14-08-1303

Chung, J., Park, J., Lai, Z. W., Lambert, T. J., Richards, R. C., Zhang, J., Walther, T. C., & Farese, R. V. (2023). The Troyer syndrome protein spartin mediates selective autophagy of lipid droplets. Nature Cell Biology, 25(8), 1101–1110. 10.1038/s41556-023-01178-w

Cinato, M., Andersson, L., Miljanovic, A., Laudette, M., Kunduzova, O., Borén, J., & Levin, M. C. (2024). Role of Perilipins in Oxidative Stress—Implications for Cardiovascular Disease. Antioxidants, 13(2), 209. https://www.mdpi.com/2076-3921/13/2/209

De Duve, C., Pressman, B. C., Gianetto, R., Wattiaux, R., & Appelmans, F. (1955). Tissue fractionation studies. 6. Intracellular distribution patterns of enzymes in rat-liver tissue. Biochem J, 60(4), 604-617. 10.1042/bj0600604

Diep, D. T. V., Collado, J., Hugenroth, M., Fausten, R. M., Percifull, L., Wälte, M., Schuberth, C., Schmidt, O., Fernández-Busnadiego, R., & Bohnert, M. (2024). A metabolically controlled contact site between vacuoles and lipid droplets in yeast. Dev Cell, 59(6), 740–758.e710. 10.1016/j.devcel.2024.01.016

Fairman, G., & Ouimet, M. (2022). Lipophagy pathways in yeast are controlled by their distinct modes of induction. Yeast, 39(8), 429–439. 10.1002/yea.3705

Farese, R. V., Jr., & Walther, T. C. (2009). Lipid droplets finally get a little R-E-S-P-E-C-T. Cell, 139(5), 855–860. 10.1016/j.cell.2009.11.005

Gao, Q., Binns, D. D., Kinch, L. N., Grishin, N. V., Ortiz, N., Chen, X., & Goodman, J. M. (2017). Pet10p is a yeast perilipin that stabilizes lipid droplets and promotes their assembly. J Cell Biol, 216(10), 3199–3217. 10.1083/jcb.201610013

Garcia, E. J., Liao, P. C., Tan, G., Vevea, J. D., Sing, C. N., Tsang, C. A., McCaffery, J. M., Boldogh, I. R., & Pon, L. A. (2021). Membrane dynamics and protein targets of lipid droplet microautophagy during ER stress-induced proteostasis in the budding yeast, Saccharomyces cerevisiae. Autophagy, 17(9), 2363–2383. 10.1080/15548627.2020.1826691

González Montoro, A., Auffarth, K., Hönscher, C., Bohnert, M., Becker, T., Warscheid, B., Reggiori, F., van der Laan, M., Fröhlich, F., & Ungermann, C. (2018). Vps39 Interacts with Tom40 to Establish One of Two Functionally Distinct Vacuole-Mitochondria Contact Sites. Dev Cell, 45(5), 621–636.e627. 10.1016/j.devcel.2018.05.011

Goodman, J. M. (2021). The importance of microlipophagy in liver. Proc Natl Acad Sci U S A, 118(2). 10.1073/pnas.2024058118

Green, D. R., Galluzzi, L., & Kroemer, G. (2011). Mitochondria and the autophagy-inflammation-cell death axis in organismal aging. Science, 333(6046), 1109–1112. 10.1126/science.1201940

Haas, A., Scheglmann, D., Lazar, T., Gallwitz, D., & Wickner, W. (1995). The GTPase Ypt7p of Saccharomyces cerevisiae is required on both partner vacuoles for the homotypic fusion step of vacuole inheritance. Embo j, 14(21), 5258–5270. 10.1002/j.1460-2075.1995.tb00210.x

Hammoudeh, N., Soukkarieh, C., Murphy, D. J., & Hanano, A. (2023). Mammalian lipid droplets: structural, pathological, immunological and anti-toxicological roles. Progress in Lipid Research, 91, 101233. 10.1016/j.plipres.2023.101233

Hardie, D. G., & Carling, D. (1997). The AMP-Activated Protein Kinase. European Journal of Biochemistry, 246(2), 259–273. 10.1111/j.1432-1033.1997.00259.x

Ishii, A., Kurokawa, K., Hotta, M., Yoshizaki, S., Kurita, M., Koyama, A., Nakano, A., & Kimura, Y. (2019). Role of Atg8 in the regulation of vacuolar membrane invagination. Sci Rep, 9(1), 14828. 10.1038/s41598-019-51254-1

Ivanovska, I. L., Tobin, M. P., Bai, T., Dooling, L. J., & Discher, D. E. (2023). Small lipid droplets are rigid enough to indent a nucleus, dilute the lamina, and cause rupture. J Cell Biol, 222(8). 10.1083/jcb.202208123

Jandrositz, A., Petschnigg, J., Zimmermann, R., Natter, K., Scholze, H., Hermetter, A., Kohlwein, S. D., & Leber, R. (2005). The lipid droplet enzyme Tgl1p hydrolyzes both steryl esters and triglycerides in the yeast, Saccharomyces cerevisiae. Biochim Biophys Acta, 1735(1), 50–58. 10.1016/j.bbalip.2005.04.005

Jensen-Pergakes, K., Guo, Z., Giattina, M., Sturley, S. L., & Bard, M. (2001). Transcriptional regulation of the two sterol esterification genes in the yeast Saccharomyces cerevisiae. J Bacteriol, 183(17), 4950–4957. 10.1128/jb.183.17.4950-4957.2001

Jones, E. W., Zubenko, G. S., & Parker, R. R. (1982). PEP4 gene function is required for expression of several vacuolar hydrolases in Saccharomyces cerevisiae. Genetics, 102(4), 665–677. 10.1093/genetics/102.4.665

Kagohashi, Y., Sasaki, M., May, A. I., Kawamata, T., & Ohsumi, Y. (2023). The mechanism of Atg15-mediated membrane disruption in autophagy. J Cell Biol, 222(12). 10.1083/jcb.202306120

Kang, N., Jinling, T., Sisi, Y., Leiying, L., & and Gao, Q. (2024). General autophagy-dependent and -independent lipophagic processes collaborate to regulate the overall level of lipophagy in yeast. Autophagy, 20(7), 1523–1536. 10.1080/15548627.2024.2325297

Klein, I., Klug, L., Schmidt, C., Zandl, M., Korber, M., Daum, G., & Athenstaedt, K. (2016). Regulation of the yeast triacylglycerol lipases Tgl4p and Tgl5p by the presence/absence of nonpolar lipids. Mol Biol Cell, 27(13), 2014–2024. 10.1091/mbc.E15-09-0633

Kurat, C. F., Natter, K., Petschnigg, J., Wolinski, H., Scheuringer, K., Scholz, H., Zimmermann, R., Leber, R., Zechner, R., & Kohlwein, S. D. (2006). Obese yeast: triglyceride lipolysis is functionally conserved from mammals to yeast. J Biol Chem, 281(1), 491–500. 10.1074/jbc.M508414200

Lei, Y., Zhang, X., Xu, Q., Liu, S., Li, C., Jiang, H., Lin, H., Kong, E., Liu, J., Qi, S., Li, H., Xu, W., & Lu, K. (2021). Autophagic elimination of ribosomes during spermiogenesis provides energy for flagellar motility. Dev Cell, 56(16), 2313–2328.e2317. 10.1016/j.devcel.2021.07.015

Li, D., Song, J. Z., Li, H., Shan, M. H., Liang, Y., Zhu, J., & Xie, Z. (2015). Storage lipid synthesis is necessary for autophagy induced by nitrogen starvation. FEBS Lett, 589(2), 269–276. 10.1016/j.febslet.2014.11.050

Londos, C., Sztalryd, C., Tansey, J. T., & Kimmel, A. R. (2005). Role of PAT proteins in lipid metabolism. Biochimie, 87(1), 45–49. 10.1016/j.biochi.2004.12.010

Menon, D., Bhapkar, A., Manchandia, B., Charak, G., Rathore, S., Jha, R. M., Nahak, A., Mondal, M., Omrane, M., Bhaskar, A. K., Thukral, L., Thiam, A. R., & Gandotra, S. (2023). ARL8B mediates lipid droplet contact and delivery to lysosomes for lipid remobilization. Cell Rep, 42(10), 113203. 10.1016/j.celrep.2023.113203

Mochida, K., Oikawa, Y., Kimura, Y., Kirisako, H., Hirano, H., Ohsumi, Y., & Nakatogawa, H. (2015). Receptor-mediated selective autophagy degrades the endoplasmic reticulum and the nucleus. Nature, 522(7556), 359–362. 10.1038/nature14506

Moir, R. D., Gross, D. A., Silver, D. L., & Willis, I. M. (2012). SCS3 and YFT2 Link Transcription of Phospholipid Biosynthetic Genes to ER Stress and the UPR. PLoS Genetics, 8.

Munzel, L., Neumann, P., Otto, F. B., Krick, R., Metje-Sprink, J., Kroppen, B., Karedla, N., Enderlein, J., Meinecke, M., Ficner, R., & Thumm, M. (2021). Atg21 organizes Atg8 lipidation at the contact of the vacuole with the phagophore. Autophagy, 17(6), 1458–1478. 10.1080/15548627.2020.1766332

Nakatogawa, H., Suzuki, K., Kamada, Y., & Ohsumi, Y. (2009). Dynamics and diversity in autophagy mechanisms: lessons from yeast. Nat Rev Mol Cell Biol, 10(7), 458–467. 10.1038/nrm2708

Oelkers, P., Cromley, D., Padamsee, M., Billheimer, J. T., & Sturley, S. L. (2002). The DGA1 gene determines a second triglyceride synthetic pathway in yeast. J Biol Chem, 277(11), 8877–8881. 10.1074/jbc.M111646200

Oelkers, P., Tinkelenberg, A., Erdeniz, N., Cromley, D., Billheimer, J. T., & Sturley, S. L. (2000). A lecithin cholesterol acyltransferase-like gene mediates diacylglycerol esterification in yeast. J Biol Chem, 275(21), 15609–15612. 10.1074/jbc.C000144200

Olzmann, J. A., & Carvalho, P. (2019). Dynamics and functions of lipid droplets. Nat Rev Mol Cell Biol, 20(3), 137–155. 10.1038/s41580-018-0085-z

Orlicky, D. J., Degala, G., Greenwood, C., Bales, E. S., Russell, T. D., & McManaman, J. L. (2008). Multiple functions encoded by the N-terminal PAT domain of adipophilin. J Cell Sci, 121(Pt 17), 2921–2929. 10.1242/jcs.026153

Paar, M., Jüngst, C., Steiner, N. A., Magnes, C., Sinner, F., Kolb, D., Lass, A., Zimmermann, R., Zumbusch, A., Kohlwein, S. D., & Wolinski, H. (2012). Remodeling of lipid droplets during lipolysis and growth in adipocytes. J Biol Chem, 287(14), 11164–11173. 10.1074/jbc.M111.316794

Pressly, J. D., Gurumani, M. Z., Varona Santos, J. T., Fornoni, A., Merscher, S., & Al-Ali, H. (2022). Adaptive and maladaptive roles of lipid droplets in health and disease. Am J Physiol Cell Physiol, 322(3), C468–c481. 10.1152/ajpcell.00239.2021

Pu, M., Zheng, W., Zhang, H., Wan, W., Peng, C., Chen, X., Liu, X., Xu, Z., Zhou, T., Sun, Q., Neculai, D., & Liu, W. (2022). ORP8 acts as a lipophagy receptor to mediate lipid droplet turnover. Protein & Cell, 14(9), 653–667. 10.1093/procel/pwac063

Ranganathan, P. R., Narayanan, A. K., Nawada, N., Rao, M. J., Reju, K. S., Priya, S. C., Gujarathi, T., Manjithaya, R., & Dodaghatta Krishna, V. R. (2022). Diacylglycerol kinase alleviates autophagic degradation of the endoplasmic reticulum in SPT10-deficient yeast to enhance triterpene biosynthesis. FEBS Lett, 596(14), 1778–1794. 10.1002/1873-3468.14418

Rao, M. J., & Goodman, J. M. (2021). Seipin: harvesting fat and keeping adipocytes healthy. Trends Cell Biol, 31(11), 912–923. 10.1016/j.tcb.2021.06.003

Rao, M. J., Srinivasan, M., & Rajasekharan, R. (2018). Cell size is regulated by phospholipids and not by storage lipids in Saccharomyces cerevisiae. Curr Genet, 64(5), 1071–1087. 10.1007/s00294-018-0821-0

Rowe, E. R., Mimmack, M. L., Barbosa, A. D., Haider, A., Isaac, I., Ouberai, M. M., Thiam, A. R., Patel, S., Saudek, V., Siniossoglou, S., & Savage, D. B. (2016). Conserved Amphipathic Helices Mediate Lipid Droplet Targeting of Perilipins 1-3. J Biol Chem, 291(13), 6664–6678. 10.1074/jbc.M115.691048

Sandager, L., Gustavsson, M. H., Ståhl, U., Dahlqvist, A., Wiberg, E., Banas, A., Lenman, M., Ronne, H., & Stymne, S. (2002). Storage lipid synthesis is non-essential in yeast. J Biol Chem, 277(8), 6478–6482. 10.1074/jbc.M109109200

Schott, M. B., Rozeveld, C. N., Weller, S. G., & McNiven, M. A. (2022). Lipophagy at a glance. J Cell Sci, 135(5). 10.1242/jcs.259402

Schott, M. B., Weller, S. G., Schulze, R. J., Krueger, E. W., Drizyte-Miller, K., Casey, C. A., & McNiven, M. A. (2019). Lipid droplet size directs lipolysis and lipophagy catabolism in hepatocytes. J Cell Biol, 218(10), 3320–3335. 10.1083/jcb.201803153

Schuck, S., Gallagher, C. M., & Walter, P. (2014). ER-phagy mediates selective degradation of endoplasmic reticulum independently of the core autophagy machinery. J Cell Sci, 127(Pt 18), 4078–4088. 10.1242/jcs.154716

Schulze, R. J., Krueger, E. W., Weller, S. G., Johnson, K. M., Casey, C. A., Schott, M. B., & McNiven, M. A. (2020). Direct lysosome-based autophagy of lipid droplets in hepatocytes. Proc Natl Acad Sci U S A, 117(51), 32443–32452. 10.1073/pnas.2011442117

Seo, A. Y., Lau, P. W., Feliciano, D., Sengupta, P., Gros, M. A. L., Cinquin, B., Larabell, C. A., & Lippincott-Schwartz, J. (2017). AMPK and vacuole-associated Atg14p orchestrate μ-lipophagy for energy production and long-term survival under glucose starvation. Elife, 6. 10.7554/eLife.21690

Shvarev, D., Schoppe, J., König, C., Perz, A., Füllbrunn, N., Kiontke, S., Langemeyer, L., Januliene, D., Schnelle, K., Kümmel, D., Fröhlich, F., Moeller, A., & Ungermann, C. (2022). Structure of the HOPS tethering complex, a lysosomal membrane fusion machinery. Elife, 11, e80901. 10.7554/eLife.80901

Singh, R., Kaushik, S., Wang, Y., Xiang, Y., Novak, I., Komatsu, M., Tanaka, K., Cuervo, A. M., & Czaja, M. J. (2009). Autophagy regulates lipid metabolism. Nature, 458(7242), 1131–1135. 10.1038/nature07976

Sorger, D., Athenstaedt, K., Hrastnik, C., & Daum, G. (2004). A yeast strain lacking lipid particles bears a defect in ergosterol formation. J Biol Chem, 279(30), 31190–31196. 10.1074/jbc.M403251200

Speer, N. O., Braun, R. J., Reynolds, E. G., Brudnicka, A., Swanson, J. M. J., & Henne, W. M. (2023). Tld1 is a regulator of triglyceride lipolysis that demarcates a lipid droplet subpopulation. Journal of Cell Biology, 223(1). 10.1083/jcb.202303026

Sturgeon, C. M., Robinson, M. R., Penton, M. C., Clemmer, D. C., Trujillo, M. A., Khawaja, A. U., & Segarra, V. A. (2019). Kinetic assay of starvation sensitivity in yeast autophagy mutants allows for the identification of intermediary phenotypes. BMC Research Notes, 12(1), 505. 10.1186/s13104-019-4545-0

van Zutphen, T., Todde, V., de Boer, R., Kreim, M., Hofbauer, H. F., Wolinski, H., Veenhuis, M., van der Klei, I. J., & Kohlwein, S. D. (2014). Lipid droplet autophagy in the yeast Saccharomyces cerevisiae. Mol Biol Cell, 25(2), 290–301. 10.1091/mbc.E13-08-0448

Vevea, J. D., Garcia, E. J., Chan, R. B., Zhou, B., Schultz, M., Di Paolo, G., McCaffery, J. M., & Pon, L. A. (2015). Role for Lipid Droplet Biogenesis and Microlipophagy in Adaptation to Lipid Imbalance in Yeast. Dev Cell, 35(5), 584–599. 10.1016/j.devcel.2015.11.010

Wada, Y., Ohsumi, Y., & Anraku, Y. (1992). Genes for directing vacuolar morphogenesis in Saccharomyces cerevisiae. I. Isolation and characterization of two classes of vam mutants. J Biol Chem, 267(26), 18665–18670.

Wang, F., Gómez-Sintes, R., & Boya, P. (2018). Lysosomal membrane permeabilization and cell death. Traffic, 19(12), 918–931. 10.1111/tra.12613

Watanabe, Y., Iwasaki, Y., Sasaki, K., Motono, C., Imai, K., & Suzuki, K. (2023). Atg15 is a vacuolar phospholipase that disintegrates organelle membranes. Cell Reports, 42(12), 113567. 10.1016/j.celrep.2023.113567

Wichmann, H., Hengst, L., & Gallwitz, D. (1992). Endocytosis in yeast: evidence for the involvement of a small GTP-binding protein (Ypt7p). Cell, 71(7), 1131–1142. 10.1016/s0092-8674(05)80062-5

Wickner, W., & Rizo, J. (2017). A cascade of multiple proteins and lipids catalyzes membrane fusion. Mol Biol Cell, 28(6), 707–711. 10.1091/mbc.E16-07-0517

Yamagata, M., Obara, K., & Kihara, A. (2011). Sphingolipid synthesis is involved in autophagy in Saccharomyces cerevisiae. Biochem Biophys Res Commun, 410(4), 786–791. 10.1016/j.bbrc.2011.06.061

Yamaguchi, T. (2007). [PAT family: lipid droplet-associated proteins that regulate fat storage and lipolysis]. Seikagaku, 79(2), 162–166.

Yen, W. L., & Klionsky, D. J. (2008). How to live long and prosper: autophagy, mitochondria, and aging. Physiology (Bethesda), 23, 248–262. 10.1152/physiol.00013.2008

Yuan, Z., Cai, K., Li, J., Chen, R., Zhang, F., Tan, X., Jiu, Y., Chang, H., Hu, B., Zhang, W., & Ding, B. (2024). ATG14 targets lipid droplets and acts as an autophagic receptor for syntaxin18-regulated lipid droplet turnover. Nature Communications, 15(1), 631. 10.1038/s41467-024-44978-w

Zechner, R., Madeo, F., & Kratky, D. (2017). Cytosolic lipolysis and lipophagy: two sides of the same coin. Nature Reviews Molecular Cell Biology, 18(11), 671–684. 10.1038/nrm.2017.76

Zechner, R., Zimmermann, R., Eichmann, T. O., Kohlwein, S. D., Haemmerle, G., Lass, A., & Madeo, F. (2012). FAT SIGNALS--lipases and lipolysis in lipid metabolism and signaling. Cell Metab, 15(3), 279–291. 10.1016/j.cmet.2011.12.018

Zhang, A., Meng, Y., Li, Q., & Liang, Y. (2020). The endosomal sorting complex required for transport complex negatively regulates Erg6 degradation under specific glucose restriction conditions. Traffic, 21(7), 488–502. 10.1111/tra.12732

Zweytick, D., Leitner, E., Kohlwein, S. D., Yu, C., Rothblatt, J., & Daum, G. (2000). Contribution of Are1p and Are2p to steryl ester synthesis in the yeast Saccharomyces cerevisiae. Eur J Biochem, 267(4), 1075–1082. 10.1046/j.1432-1327.2000.01103.x

