## Supplementary figures and images for "Pln1 Mediates Lipid Droplet-Vacuole Tethering During Microlipophagy in *Saccharomyces cerevisiae*"

### Fig. S1

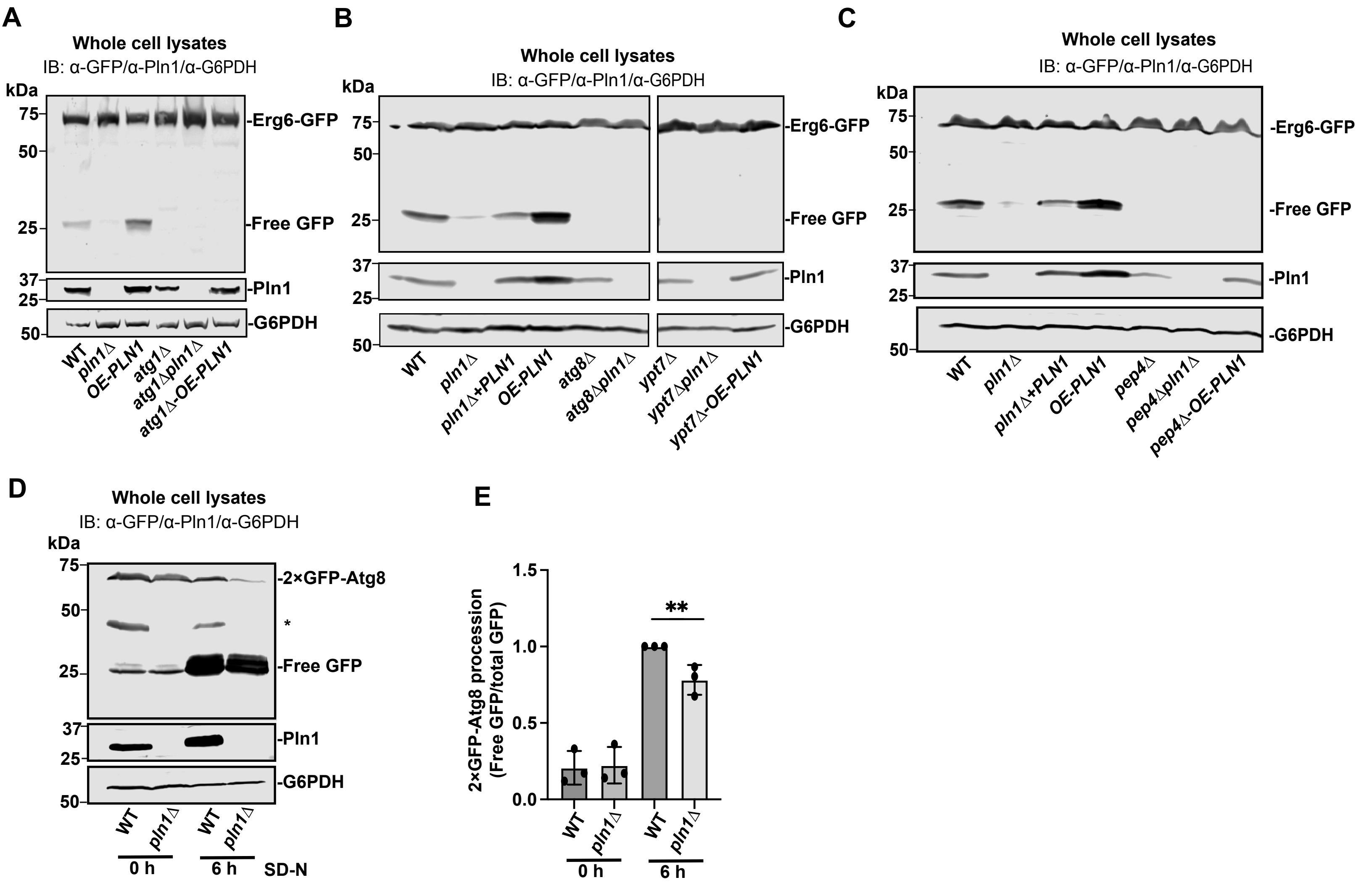

### Fig. S2

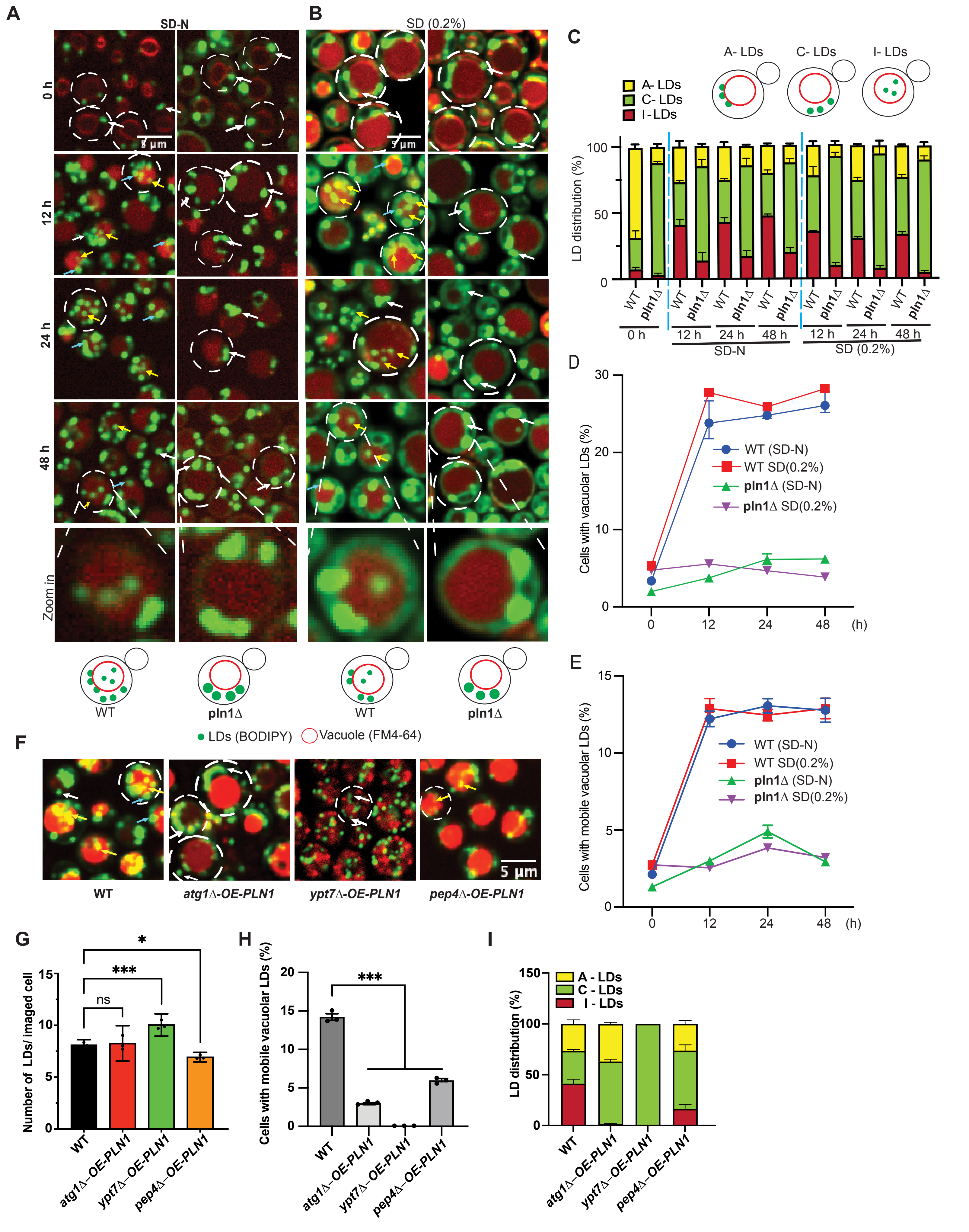

### Fig. S3

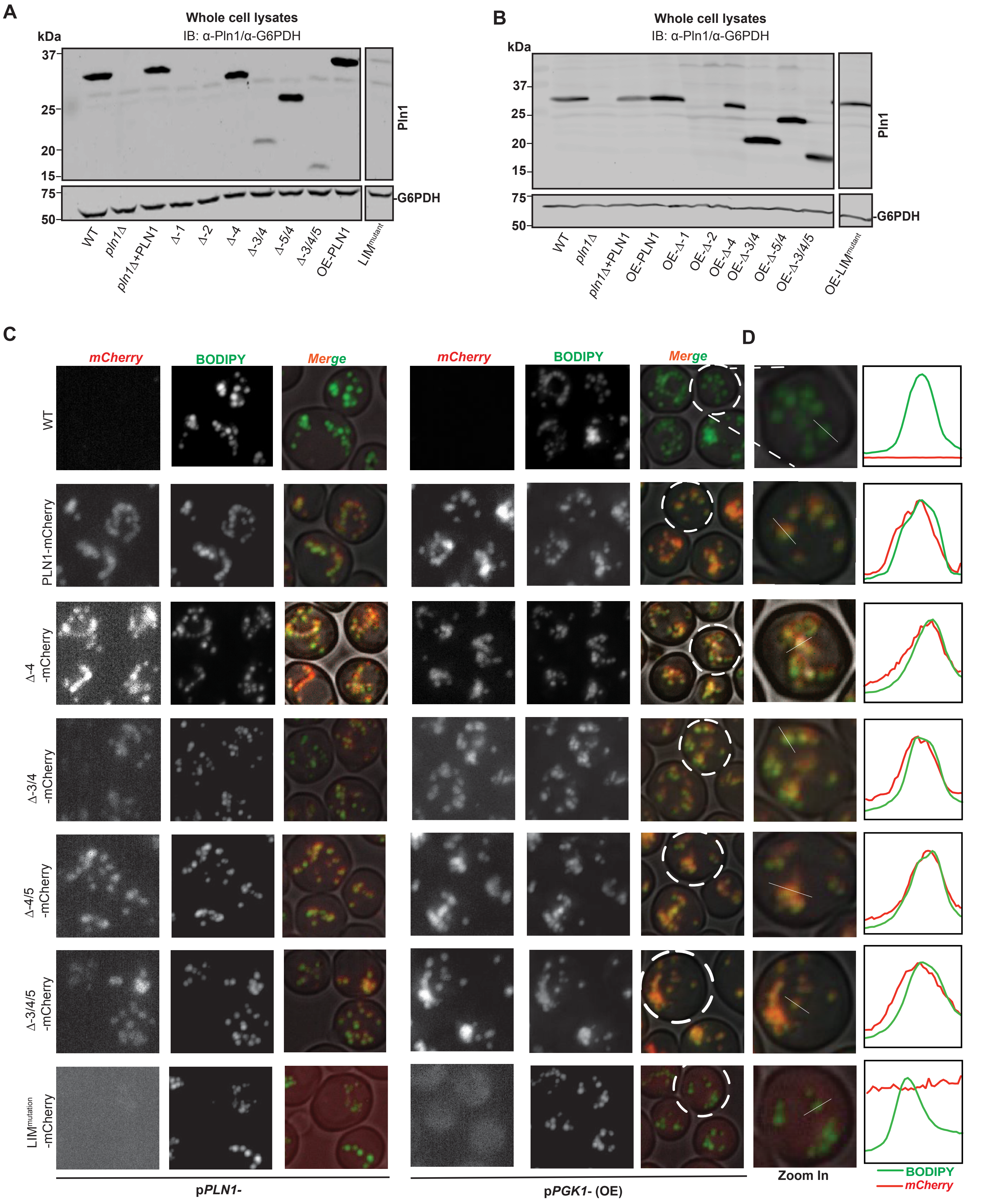

### Fig. S4

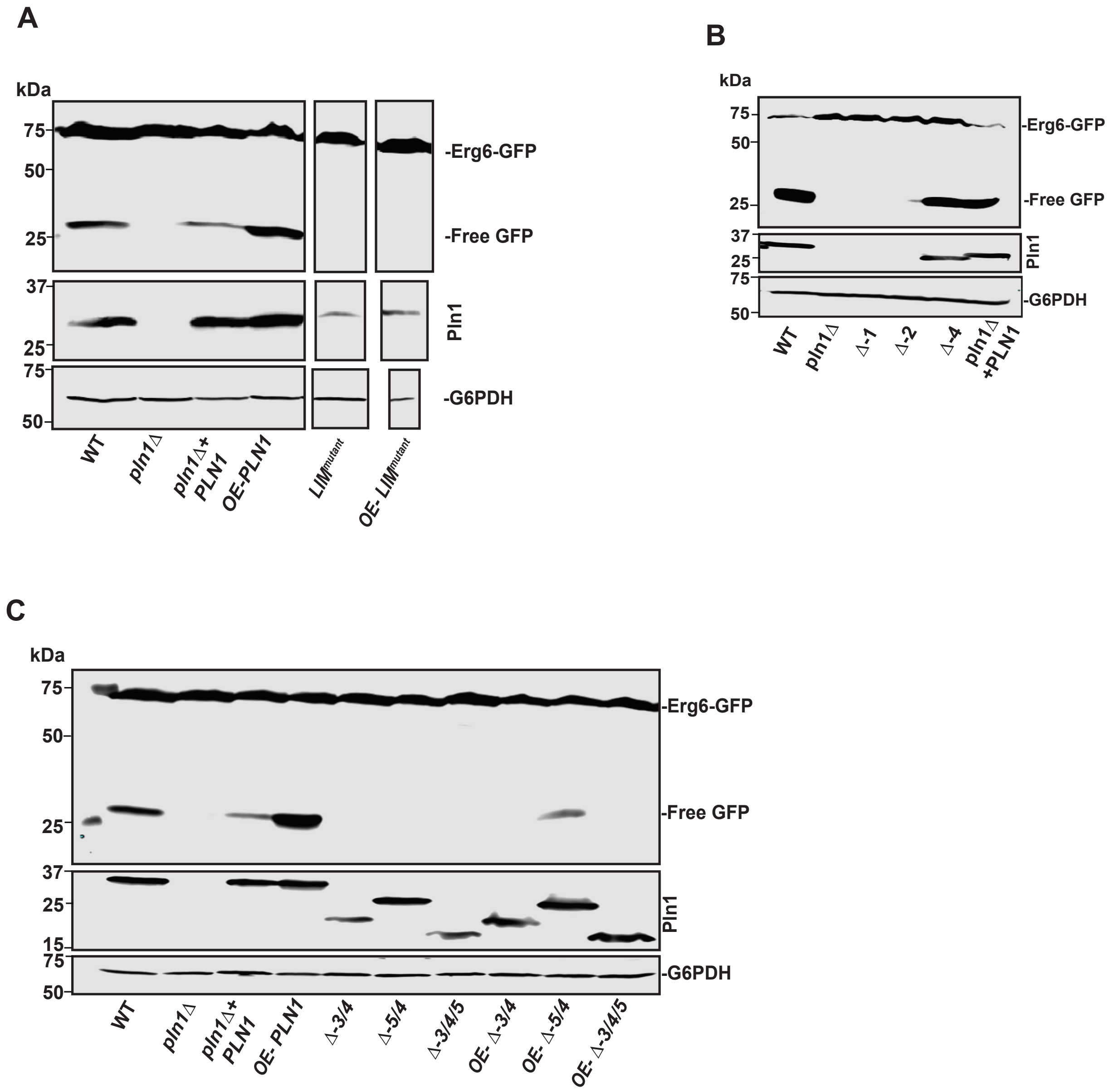

### Fig. S5

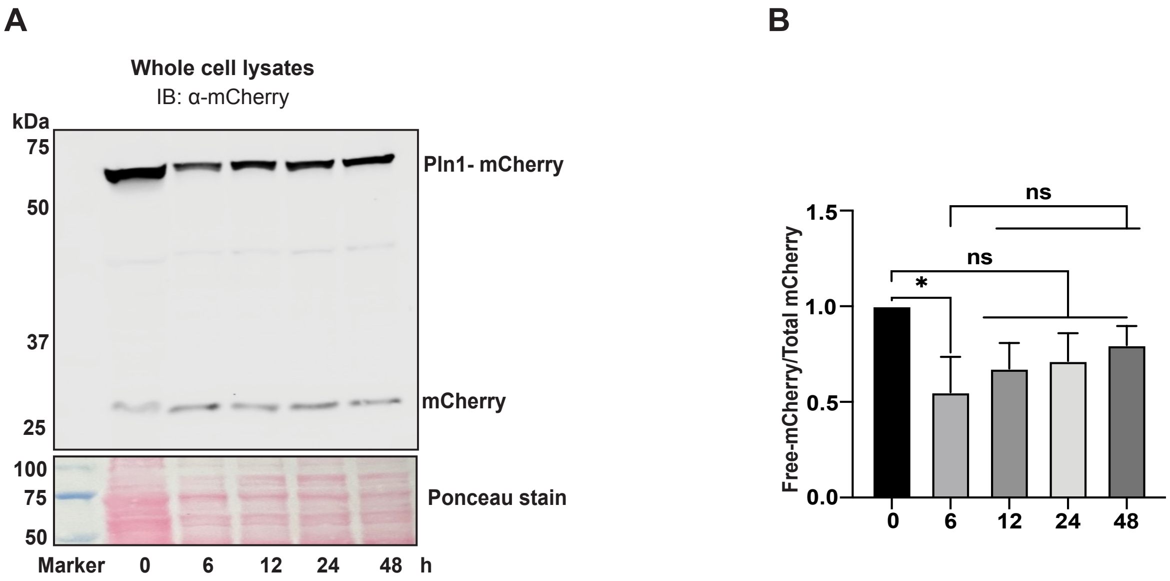

### Fig. S6

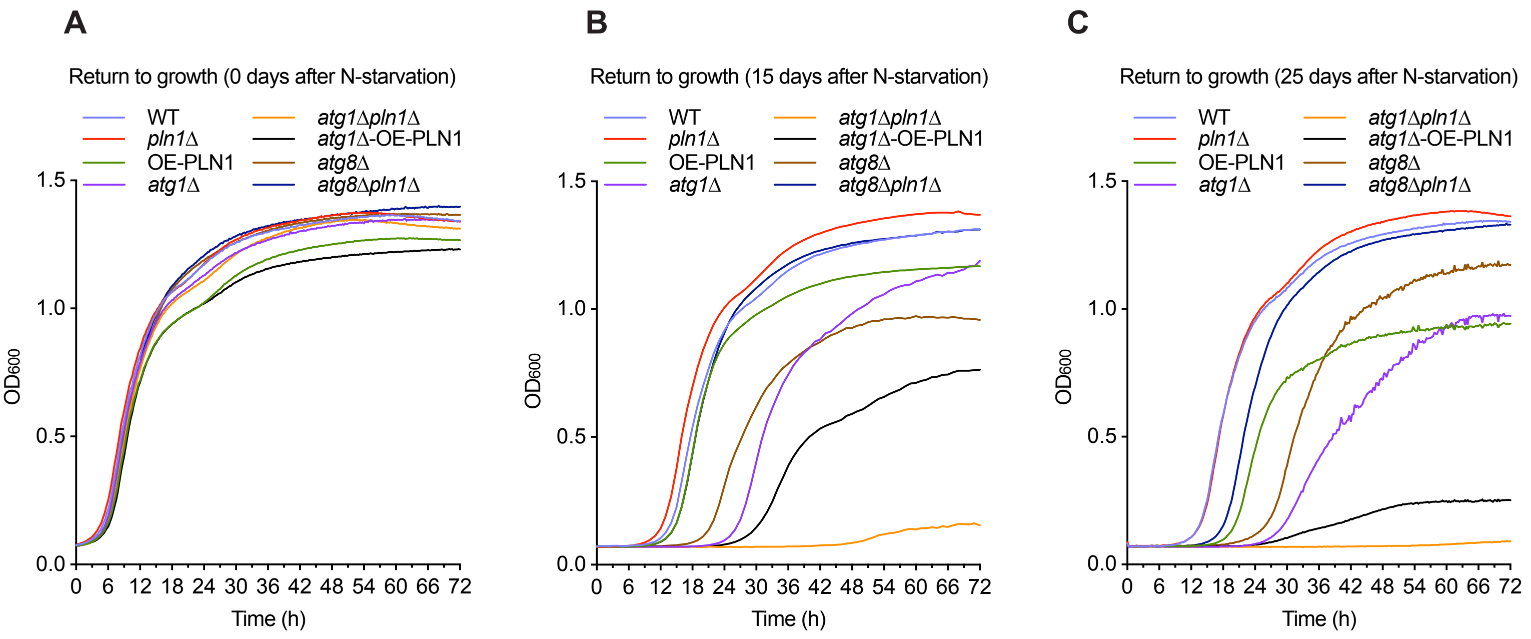
